# Systematic Evaluation of High-Throughput PBK Modelling Strategies for the Prediction of Intravenous and Oral Pharmacokinetics in Humans

**DOI:** 10.1101/2024.03.20.585001

**Authors:** René Geci, Domenico Gadaleta, Marina García de Lomana, Rita Ortega-Vallbona, Erika Colombo, Eva Serrano-Candelas, Alicia Paini, Lars Kuepfer, Stephan Schaller

## Abstract

Physiologically based kinetic (PBK) modelling offers a mechanistic basis for predicting the pharmaco-/toxicokinetics of compounds and thereby provides critical information for integrating toxicity and exposure data to replace animal testing with *in vitro* or *in silico* methods. However, traditional PBK modelling depends on animal and human data, which limits its usefulness for Non-Animal Methods. To address this limitation, High-throughput PBK modelling aims to rely exclusively on *in vitro* and *in silico* data for model generation. Here, we evaluate a variety of *in silico* tools and different strategies to parameterise PBK models with input values from various sources in a high-throughput manner. We gather 2000+ publicly available human *in vivo* concentration-time profiles of 200+ compounds (IV and oral administration), as well as *in silico*, *in vitro* and *in vivo* determined compound-specific parameters required for the PBK modelling of these compounds. Then, we systematically evaluate all possible PBK model parametrisation strategies in PK-Sim and quantify their prediction accuracy against the collected *in vivo* concentration-time profiles. Our results show that even simple, generic High-throughput PBK modelling can provide accurate predictions of the pharmacokinetics of most compounds (87% of Cmax and 84% of AUC within 10-fold). Nevertheless, we also observe major differences in prediction accuracies between the different parameterisation strategies, as well as between different compounds. Finally, we outline a strategy for High-throughput PBK modelling that relies exclusively on freely available tools. Our findings contribute to a more robust understanding of the reliability of High-throughput PBK modelling, which is essential to establish the confidence necessary for its utilisation in Next-Generation Risk Assessment.

## Introduction

Pharmacokinetics (PK) and toxicokinetics (TK), the study of the distribution of drugs or chemicals within the body over time, are fundamental for understanding the desired or undesired effects of compounds on human health. Physiologically based pharmacokinetic (PBPK) or physiologically based toxicokinetic (PBTK) models, hereafter summarised as physiologically based kinetic (PBK) models, are a well-established computational method for simulating the PK of drugs, or TK of chemicals (Jones and Rowland-Yeo 2013). PBK models incorporate anatomical and physiological knowledge about the body, such as tissue compositions and blood flow rates, as well as compound-specific properties like the lipophilicity or solubility of compounds, to simulate the absorption, distribution, metabolism, and excretion (ADME) processes that determine compound concentrations in the body. This makes PBK models powerful tools that enable a mechanism-based prediction of compounds’ concentration-time profiles in plasma and body tissues, even those otherwise inaccessible for direct sampling.

In the pharmaceutical industry, PBK models play a crucial role in drug discovery and development, cross-species extrapolation, the guiding of dosing regimens, extrapolation to special populations, and the prediction of potential drug-drug interactions (Thiel et al. 2015; Krstevska et al. 2022). Their use reduces drug failure rates and optimises trial protocols (Edginton et al. 2008; Lin et al. 2022). Further, PBK models are now also used outside the pharmaceutical field from which they originated. In toxicology, PBK models have become more widely adopted and aid in interpreting human biomonitoring data (Clewell et al. 2008), in the extrapolation from *in vitro* to *in vivo* data (Bouvier d’Yvoire et al. 2007; Blaauboer 2010; Yoon et al. 2012) and in chemical safety assessment (Paini et al. 2019a; Paini et al. 2021). While pharmacology and toxicology have distinct objectives, both aim to understand the concentrations of compounds in the body, making PBK models a valuable tool in both fields.

Traditionally, generating PBK models is an iterative, time- and labour-intensive process. It is performed on a compound-by-compound basis and requires animal and human data for model parameterisation and validation. This makes the use of PBK models low-throughput and limits their usefulness for chemical risk assessment. Further, there is now a general desire in both the pharmaceutical and toxicological field to reduce the use of animal data to comply with the 3R principles (Törnqvist et al. 2014; Stokes 2015). For these reasons, there has been increasing interest to explore PBK modelling exclusively based on *in vitro* and *in silico* methods, as these methods have a greater potential to be applied in high-throughput and promise to reduce dependence on animal data. This new approach to PBK modelling is sometimes called High-throughput PBK (HT-PBK) modelling (Pearce et al. 2017; Breen et al. 2021; Naga et al. 2022; Khalidi et al. 2022) or Next-Generation PBK modelling (Paini et al. 2019b; Punt et al. 2022a), either emphasising its rapid nature or that it does not rely on animal data.

The shift towards basing PBK modelling on *in silico* methods is further supported by recent advances in the fields of machine learning (ML) and artificial intelligence (AI) (Bender and Cortés-Ciriano 2021). ML and AI technologies are now increasingly being explored to provide rapid PK predictions without the need for new animal data. Sometimes, this is pursued by using AI/ML methods to directly predict summary PK/TK parameters, like maximum concentration (Cmax) or area under the curve (AUC) values (Miljković et al. 2021; Fagerholm et al. 2021; Li et al. 2023). Other times, it is done by predicting mechanistically relevant compound properties, like the lipophilicity, solubility or clearance of compounds, which can then be used as inputs for making PK predictions using mechanistic models (Danishuddin et al. 2022; Pillai et al. 2022; Fagerholm et al. 2023; Mavroudis et al. 2023; Führer et al. 2024). Until now, many *in silico*-based PK prediction efforts have focused on predicting rodent data (Schneckener et al. 2019; Kamiya et al. 2021; Naga et al. 2022; Punt et al. 2022b; Obrezanova et al. 2022; Mavroudis et al. 2023; Handa et al. 2023), presumably due to its greater availability than human data. While valuable, the ultimate goal in pharmacology and toxicology is to yield human-relevant conclusions. In those instances where such approaches were developed for predicting human data, evaluations were usually performed against relatively small datasets, typically not exceeding a few dozen compounds (Punt et al. 2022a; Li et al. 2023; Fagerholm et al. 2023). Moreover, PK prediction validations frequently relied on summary PK parameters, like Cmax or AUC values (Miljković et al. 2021). While they can provide useful insights, summary parameters also obscure prediction inaccuracies and do not fully reflect all intricacies of PBK model quality. This is why PBK models are traditionally evaluated against full concentration-time profiles instead. Furthermore, a number of *in silico* tools capable of predicting compound properties required for PBK modelling are available already (Benfenati et al. 2013; Mansouri et al. 2018; Xiong et al. 2021). However, to date, there has been no systematic evaluation of the usefulness of these existing tools, or their various combinations, for HT-PBK modelling.

The aim of this work was to evaluate strategies for the high-throughput generation of human PBK models with input parameters from already available *in silico*-based property prediction tools. To this end, we compiled a large dataset of human *in vivo* concentration-time profiles after intravenous (IV) and oral (PO) administration (210 compounds, 2235 concentration-time profiles). Then, we systematically evaluated the predictive performances of PBK models when parameterised with the different *in silico* tools, and further compared their results to *in vitro* and *in vivo* benchmark references. This allowed us to identify which of the *in silico* tools provide the best input parameters for PBK modelling, as well as to validate the overall accuracy of such HT-PBK models for predicting IV and PO PK profiles in humans.

## Material and Methods

### Retrieval of Pharmacokinetic Data

An extensive description of the PK data retrieval process can be found in the Supplementary Information (SI-1). In short, we downloaded all concentration-time data from the CvT-DB (Sayre et al. 2020), OSP Observed Data Repository (Lippert et al. 2019), and PK-DB (Grzegorzewski et al. 2021) databases. We selected data measured in healthy adult humans after single IV or PO administration, excluding studies potentially dedicated to special patient cohorts like paediatric, geriatric, or diseased populations. Further, studies were excluded when concomitant treatments that might influence compounds’ PK had occurred, as in drug-drug interaction studies or with special treatments such as grapefruit administration. Next, we manually extended this dataset by adding available literature data for compounds of relevance to the EU-funded ONTOX project (Vinken et al. 2021). We searched the literature for corresponding PK studies, for example using PKPDAI (Gonzalez Hernandez et al. 2021), and also did the same for compounds for which we had only obtained a single concentration-time profile before. From the retrieved literature, we manually extracted concentration-time profiles using WebPlotDigitizer version 4.6 (https://automeris.io/WebPlotDigitizer; Rohatgi 2022). Eventually, we had compiled a large dataset with multiple concentration-time profiles after IV and PO administration for most compounds. To ensure the consistency of the PK data, we visually checked dose-normalised plots of each compound and route and ensured that none of the PK study data were in extreme disagreement with each other. In the case of severe outliers, we manually investigated the causes of these differences, which in most cases were digitisation errors. We then either corrected such errors or, when it was unexplainable why individual studies were in disagreement with the other studies, we excluded those outliers from the dataset.

### Retrieval of Compound Property Data

To evaluate different PBK model parameterisation strategies, we retrieved or predicted the required input properties of all compounds of which we had obtained PK data. The minimal input properties required for model parameterisation in PK-Sim are a compound’s lipophilicity, pKa values, plasma or hepatic intrinsic clearance and its fraction unbound (Fu) (Kuepfer et al. 2016). Additionally, the simulation of oral administration also requires values for the solubility and intestinal permeability.

We generated lipophilicity predictions using six different *in silico* tools: LogP values by OCHEM (https://ochem.eu/home/show.do) and VEGA ALOGP (Ghose and Crippen 1987; Benfenati et al. 2013), LogD values by ADMETLab 2.0 (Xiong et al. 2021) and ADMETPredictor version 11.0.0.3 (henceforth called “SimPlus”; https://www.simulations-plus.com/software/admetpredictor), and predictions from Bayer’s inhouse models of LogD and LogMA (Göller et al. 2020). We further converted LogP or LogD values to LogMA-type values using regression relationships taken from Yun and Edginton (2013), Endo et al. (2011), and Loidl-Stahlhofen et al. (2001). pKa values were predicted using ChemAxon (Lee and Crippen 2009) and ADMETPredictor, aqueous solubility values using four *in silico* tools named ProtoPRED (https://protopred.protoqsar.com/), OPERA (Mansouri et al. 2018), ADMETPredictor, and ADMETLab 2.0. Further, Fasted State Simulated Intestinal Fluid (FaSSIF) and Fed State Simulated Intestinal Fluid (FeSSIF) solubility was also predicted using ADMETPredictor.

Intestinal permeability was predicted in the form of CACO2 values using ADMETLab 2.0, ProtoPRED and OPERA, as well as in the form of MDCK permeability using ADMETLab 2.0 and ADMETPredictor. Fu values were predicted using seven different *in silico* tools: ADMETLab 2.0, ADMETPredictor, pkCSM (Pires et al. 2015), Watanabe et al. (2018), OPERA, VEGA logK and VEGA CORAL (Toma et al. 2018). Plasma clearance values were predicted using ADMETLab 2.0, pkCSM and the ScitoVation clearance tool (https://scitovation-testing.shinyapps.io/clearancetoolgui). Hepatocyte CLint values were predicted with OPERA, ADMETPredictor and ADMET-AI (Swanson et al. 2023).

We also retrieved experimentally measured benchmark reference values of solubility, Fu, and clearance from public sources when they were available. In particular, aqueous solubility (logS) data were retrieved from Hughes et al. (2008), and the CompTox Chemicals Dashboard (Williams et al. 2017). Fu values were collected from Tonnelier et al. (2012), Yamazaki and Kanaoka (2004), Lombardo et al. (2002), Riley et al. (2005), Sohlenius-Sternbeck et al. (2010), Votano et al. (2006), Zhu et al. (2013), from the CompTox Chemicals Dashboard and the R httk package version 2.2.1 (Pearce et al. 2017). We also retrieved intrinsic hepatocyte clearance values from the R httk package version 2.2.1 (Pearce et al. 2017), as well as fitted PK-Sim intestinal permeability values from Willmann et al. (2004). Finally, we retrieved *in vivo* observed plasma clearance values from the literature, preferentially from IV studies, except for compounds for which no IV studies were available. When multiple values of a property were acquired for a single compound, the average values were used. And when experimental values of a compound were found in the previously described *in silico* tool’s training set, those values were also added to the experimental data retrieved from the aforementioned datasets.

### Software, Model Parameterisation and Performance Metrics

All PBK model simulations were performed with PK-Sim version 11.1.137 (Willmann et al. 2003) executed from R using the ospsuite package version 11.1.143 and R version 4.2.2 (R Core Team 2022). For every retrieved PK study, a corresponding PBK model simulation was performed by parameterising a generic PBK model with a) all compound-specific parameter values as provided by the different tools in the different parameterisation strategies, b) the study-specific parameters like route of administration, dose and infusion duration (IV) or formulation (PO), if provided in the PK data, and c) demographic parameters like sex, age, weight, and height of subjects, if provided in the PK data. When demographic parameters were not available, PK-Sim default settings (healthy adult male) were used for simulation.

Predicted or measured hepatic intrinsic clearance (CLint) values were scaled to *in vivo* liver clearance as

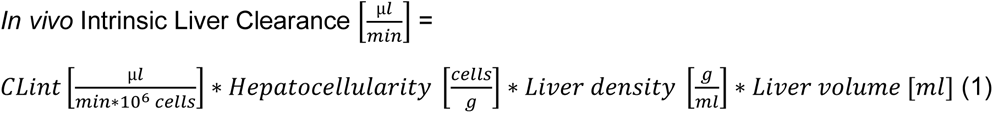

using the same values as Pearce et al. (2017) with a hepatocellularity of 1.1x10^8^ cells/g (Ito and Houston 2004), a liver density of 1.05 g/ml (Snyder WS et al. 1979) and the PK-Sim default liver volume.

For oral simulations, we also set a formulation-specific dissolution parameter (80% dissolution time of Lint80 formulation as defined in PK-Sim), as some of the oral data was not obtained by administration of drugs in solution but, for example, in the form of capsules or tablets which are not dissolved immediately upon administration. The values used for this were 25 min for “capsule” and 40 min for “tablet” formulations.

After simulating PBK models, the simulation results and the observed concentration-time data of corresponding PK studies were compared against each other and different model performance metrics were calculated to quantify PBK model quality: Log2-fold changes of predicted/observed values of Cmax, Tmax, AUC 0-last calculated using the linear up-log down method of the PKNCA package version 0.10.0 (Denney et al. 2015), as well as the percentage of datapoints within different fold ranges (1.5/2/3/5/10-fold). Further, we evaluated the goodness of the entire concentration-time profile prediction against all datapoints in the *in vivo* PK profile calculating the Relative and Absolute Log2 Errors as

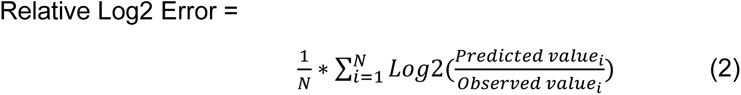

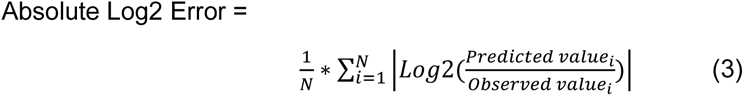

for all N datapoints in every concentration-time profile such that the Relative Log2 Error captures biases in prediction, i.e., over- or underestimates (systematic error), while the Absolute Log2 Error quantifies overall closeness of PBK model simulations to the observed concentration-time values (random error).

When we had obtained multiple concentration-time profiles of a single compound, we selected the median performance value of all studies as the representative value for the compound to ensure that individual PK study outliers would not distort our results. To later summarise the overall performance of full parameterisation strategies, we followed the same approach to integrate the results of the various compounds in our dataset, so that when we refer to the Median Log2 Error, this means the median of the Log2 Error values of all the compounds in the evaluation dataset.

## Results

### PK Data Extraction and Simulation Strategy

After the retrieval of PK data from the different databases and the literature (SI-1), we initially obtained a total of 2235 healthy human adult *in vivo* concentration-time profiles of 210 unique compounds (fig. 1A). For all compounds in this dataset, we then collected the various input data we intended to evaluate for PBK model parameterisation, for instance, lipophilicity values or predicted and measured plasma clearances. This resulted in a complete dataset of 718 IV profiles (143 compounds) and 1402 PO profiles (169 compounds) for which all compounds also had the required input data. Notable exceptions to this were *in vitro* measured intrinsic hepatic clearance values (CLint), which were taken from httk and were available for only 97 compounds, and measured solubility values which were only available for 167 compounds. The 182 compounds in the final dataset were a diverse set of small molecules with a molecular weight of less than 900 Dalton, different ionisation states at physiological pH, and mostly belonged to ECCS class 2, suggesting that metabolism is driving their clearance (fig. 2B).

**Fig. 1:**
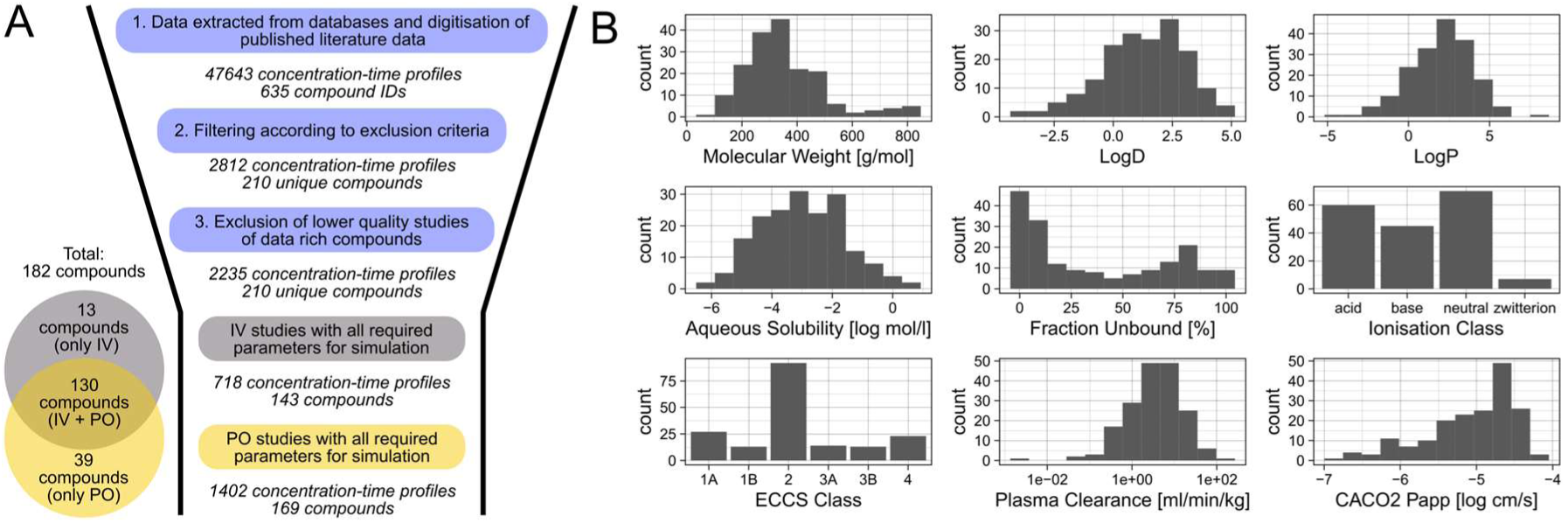
Summary statistics of the collected PK dataset and of various compound properties. A: Overview of the number of concentration-time profiles and compounds during and at the end of the PK data retrieval process. B: Physico-chemical and pharmacokinetic properties of compounds in the retrieved PK dataset. LogD, LogP, aqueous solubility and CACO2 permeability are given as the mean of values predicted in silico by the different prediction tools used in this study. Plasma clearance values are the median of in vivo values collected from the literature. Ionisation at physiological pH (7.4) was predicted using ChemAxon.

**Fig. 2:**
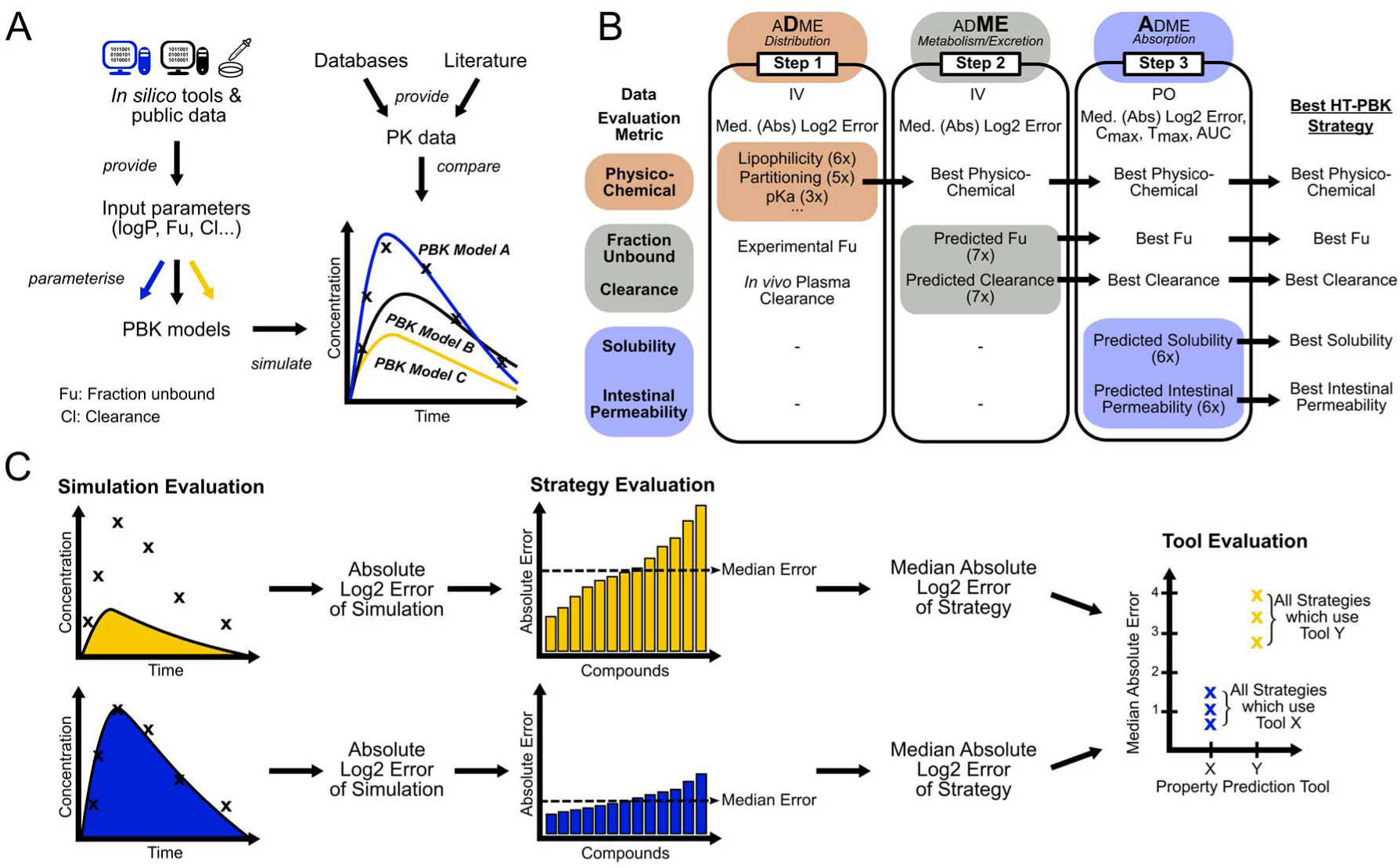
Conceptual overview of the simulation and analysis strategy. A: For each compound various PBK models were generated using all combinations of available input parameter sources. Their performance was then compared against gathered concentration-time profiles. B: The simulation and evaluation of PBK models built from the various parameterisation sources was split into three steps, each focusing on the evaluation of a separate level of ADME processes. C: Model performance was first evaluated at the level of individual simulations as the difference between model simulation and observed data. Then, for every parameterisation strategy, the median error of all compounds was used as a measure of the performance of that parameterisation strategy. Finally, comparing all performances of all strategies using the same input property prediction tools allowed benchmarking of the various tools against each other.

Evaluating the performance of all PBK models parameterised from all the different input data sources against a large number of *in vivo* PK data requires many model simulations and leads to long computation times. For this reason, we split the simulation and analysis of the HT-PBK model parameterisation strategies into three steps, to be able to systematically evaluate all relevant parameterisation decisions one-at-a-time while keeping the number of required model simulations manageable (here 15+ million).

Briefly summarised, the rationale for our simulation and analysis strategy was as follows (fig. 2). In the first step, we investigated which of the physico-chemical parameter sources performed best for predicting the passive distribution of compounds within the body. We generated, simulated, and evaluated every PBK model parameterisation strategy for all compounds of which IV data were available. To initially limit the variability, we only used *in vivo* observed plasma clearance and *in vitro* measured Fu values as high-quality benchmark reference values in the first step. In the second step, we then used the same IV data, but now only using the best physico-chemical parameter predictions as determined in the first round, and then tested various Fu and clearance prediction tools to understand which of these would result in the best PK predictions. In the third step, we finally used the oral PK data for evaluation, along with the best physico-chemical, Fu and clearance prediction sources as determined in the previous steps. Then, we systematically varied the various solubility and intestinal permeability values to evaluate how to best predict oral absorption and to assess how well the best high-throughput strategies would perform overall.

### Step 1: Evaluation of Physico-Chemical Property Predictions

In the first step, we systematically evaluated how to best set the physico-chemical PBK model parameters that determine the passive distribution of compounds within the body. In PK-Sim, these are primarily the lipophilicity of a compound, its pKa values, and the method used to predict the compound’s partitioning coefficients. The lipophilicity values of compounds were predicted with six different *in silico* tools: three LogD prediction tools (SimPlus, ADMETLab, Bayer), two LogP tools (OCHEM and VEGA), and one LogMA tool (Bayer). pKa values were predicted using ChemAxon and SimPlus, and for comparison we also tested not providing pKa values, effectively assuming that all compounds were neutral. The five tested partitioning methods available in PK-Sim were PK-Sim (Willmann et al. 2005), Schmitt (2008), Rodgers and Rowland (2006), Poulin and Theil (2002), and Berezhkovskiy (2004). To limit the variability in this first analysis, we used *in vivo* observed plasma clearance and experimentally measured Fu values as high-quality benchmark values for simulation, so that any remaining PBK model simulation inaccuracies would only be due to mispredictions of passive compound distribution alone. Then, we evaluated which physico-chemical prediction tools resulted in the best PBK model simulations by systematically testing all combinations of input parameter sources against our 718 collected concentration-time profiles after IV administration. For each simulation, we calculated Median Relative and Absolute Log2 Errors as measures of prediction bias (systematic error) and precision (random error), respectively.

Out of the tested parameters, we found the strongest factor determining PBK model accuracy was which lipophilicity values were used for PBK model building (fig. 3A). Tools that predicted LogP performed overall worse than those predicting LogD or LogMA lipophilicity values. Likewise, the performances of tools predicting the same type of lipophilicity also differed. For example, LogD values predicted by the Bayer tool worked better for PBK model parameterisation than the ones coming from ADMETLab or SimPlus (ADMETPredictor). The higher errors of the LogP tool based predictions were correlated with a general bias for underprediction of the PK data. When investigating this effect on the individual compound level (fig. 3E-J), it became apparent that for some compounds the LogP tools predicted very high lipophilicity values (> 5), which then led to a severe underprediction of those compounds’ plasma concentrations, while the same compounds’ PK was predicted reasonably well when using LogD or LogMA values for PBK model parameterisation.

**Fig. 3:**
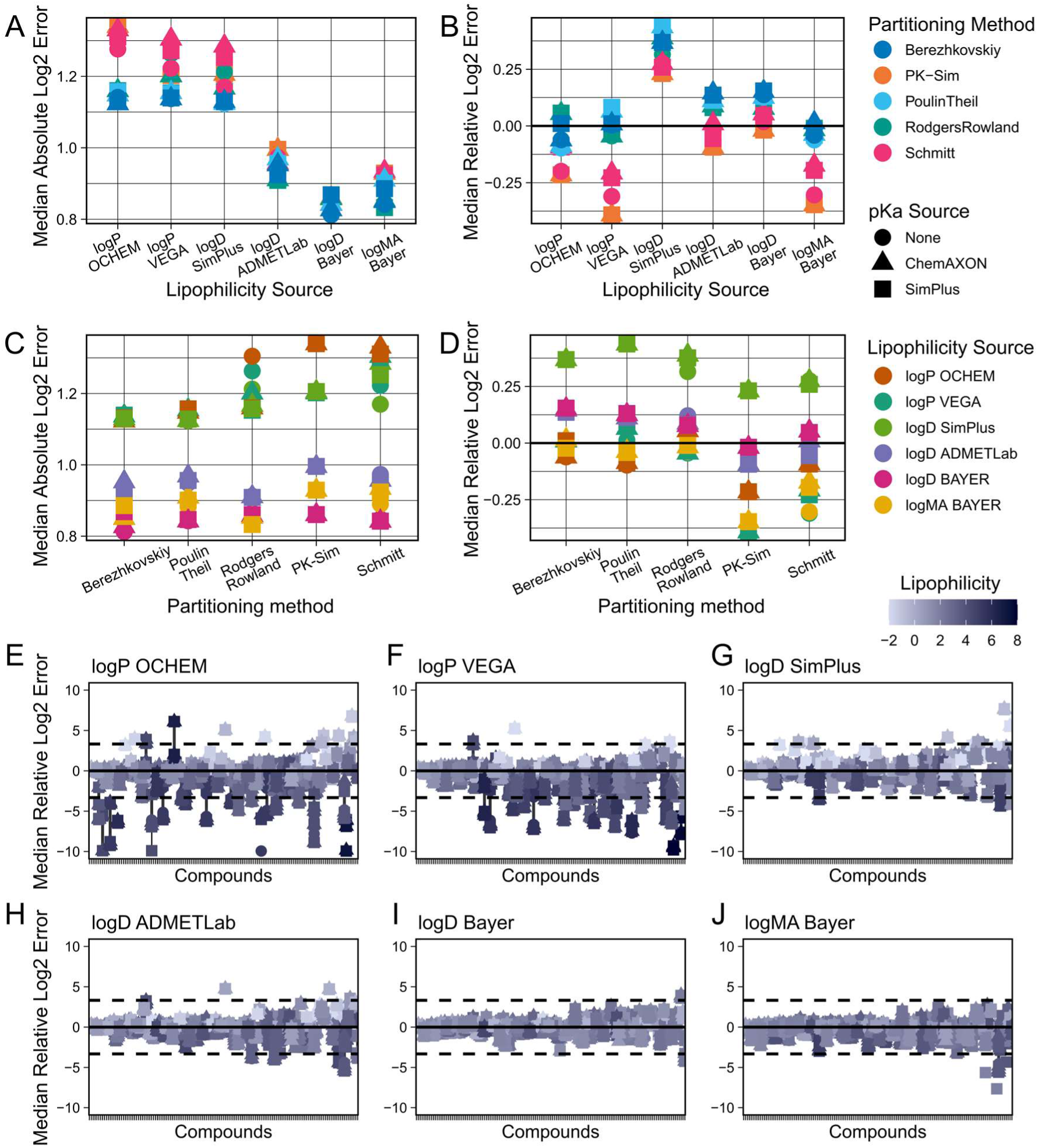
Comparison of predictive performances of different physico-chemical parameter sources (step 1). Combinations of all available PBK model parameterisation sources were evaluated against the collected IV dataset. Clearance and fraction unbound were parameterised using *in vivo* and *in vitro* benchmark reference values, respectively. The top row shows Median Absolute Log2 Errors (A) and Median Relative Log2 Errors (B) for different lipophilicity prediction methods. The middle row shows Median Absolute Log2 Errors (C) and Median Relative Log2 Errors (D) for different partitioning methods. The bottom rows (E–J) show the Relative Log2 Errors of predictions for every individual compound for the different lipophilicity prediction tools used. Dashed lines indicate 10-fold errors.

The results of the partitioning methods were less straightforward. We observed a stable hierarchy in the predictive performances of the different methods, with the Berezhkovskiy method performing best and the Schmitt method performing worst under most circumstances (fig. 3A). However, this difference in performance was only observed clearly when using lipophilicity values from the less well-performing LogP prediction tools, whereas when using the better lipophilicity values (LogD and LogMA Bayer) the difference in prediction precision between the different partitioning methods was only marginal (fig. 3C). We further investigated this at the individual compound level (SI-Fig. 1), and we observed that the methods of Poulin & Theil and of Berezhkovskiy were not generally more predictive for the majority of compounds. Rather, we found that Poulin & Theil and Berezhkovskiy were less negatively impacted by the very high lipophilicity values predicted by the LogP tools for some compounds (SI-Fig. 2). It was only their robustness to these high lipophilicity outliers, which decreased the observed performance of the other partitioning methods but not theirs, that made them appear to be superior overall (SI-Fig. 3).

For the provision of predicted pKa values, there was no strong trend observable, even though one may have expected that partitioning methods that use pKa values as input should perform better when those are provided. However, this was only consistently the case for Rodgers & Rowland partitioning, and only when it was used with LogP lipophilicity values (SI-Fig. 4). The other methods using pKa values, namely the methods of Schmitt, Poulin & Theil and Berezhkovskiy, performed sometimes better, sometimes worse, depending on which lipophilicity prediction tool was being used for simulation.

Finally, we evaluated two approaches to further improve lipophilicity predictions for PBK modelling. The first approach was that of using consensus values of the different prediction tools, e.g., for LogP, simply by taking the mean of the values predicted by the different tools. We did this with both the tools for LogP, as well as for LogD, respectively, and, interestingly, observed opposite effects (SI-Fig. 5). Averaging the LogP predictions indeed resulted in better predictivity of the PBK models than any of the individual LogP tools. For the LogD, however, averaging produced better results than the worst tool (SimPlus) but worse results than the better tools (Bayer, ADMETLab).

The second strategy we tested to improve lipophilicity predictions was to use regression equations that empirically relate LogP or LogD values to membrane affinity (LogMA). We obtained two equations for converting LogP values (Yun et al. 2014; Endo et al. 2011) and generated a comparable equation for LogD values based on the data presented in Loidl-Stahlhofen et al. (2001). But we found that only one of the three strategies, namely using the Yun et al. (2014) equation, consistently improved PK predictions, regardless of which LogP tool or partitioning method it was used with (SI-Fig. 6). Whereas the two other methods showed at best mixed results, or even worsened predictions in the case of our self-derived equation based on the Loidl-Stahlhofen et al. (2001) data. The improvement in PK predictions, achieved by converting LogP to LogMA values using the equation from Yun et al. (2014), suggests that the tested LogP prediction tools may not be inherently less accurate than the LogD tools. Rather, they may just provide a type of lipophilicity value that is less suitable for the PBK modelling of certain compound classes.

**Table 1:** Overview of different HT-PBK modelling strategies and their predictive performances.

|  |  | **Best overall**<br>(including *in vivo* and *in vitro* benchmark values) | **Best *in silico* - proprietary** | **Best *in silico* - free** |
| --- | --- | --- | --- | --- |
| **Input parameter sources** | **Lipophilicity** | Mean of Bayer LogD and LogMA | Mean of Bayer LogD and LogMA | ADMETLab 2.0: LogD |
|  | **Solubility** | SimPlus: FaSSiF | SimPlus: FaSSiF | ADMETLab 2.0: Aqueous solubility |
|  | **pKa** | ChemAxon | ChemAxon | None |
|  | **Partitioning method** | PK-Sim | PK-Sim | PK-Sim |
|  | **Fraction Unbound** | *In vitro* measured | ADMETLab 2.0 | ADMETLab 2.0 |
|  | **Clearance** | *In vivo* observed<br>Plasma clearance | ADMETLab 2.0: Plasma clearance | ADMETLab 2.0: Plasma clearance |
|  | **Intestinal permeability** | ADMETLab 2.0: MDCK | ADMETLab 2.0: MDCK | ADMETLab 2.0: MDCK |
| **Performance metrics** | **IV: Median Absolute Log2 Error** | 0.860 | 1.065 | 1.273 |
|  | **PO: Median Absolute Log2 Error** | 1.245 | 1.595 | 1.780 |
|  | **PO: Median Absolute Log2 Cmax predicted/observed** | 0.998 | 1.127 | 1.130 |
|  | **PO: Compounds with Cmax within 10/5/3/2-fold** | 86% / 78% / 69% / 50% | 87% / 78% / 65% / 44% | 89% / 71% / 62% / 42% |
|  | **PO: Underprediction of compound Cmax of more than 10/5/3/2-fold** | 1.9% / 4.3% / 8.1% / 16% | 0% / 3.1% / 8.1% / 19% | 3.1% / 12% / 16% / 25% |
|  | **PO: Median Absolute Log2 AUC predicted/observed** | 0.870 | 1.285 | 1.358 |
|  | **PO: Compounds with AUC within 10/5/3/2-fold** | 92% / 84% / 75% / 57% | 84% / 71% / 58% / 38% | 84% / 66% / 54% / 39% |

Given these results, we concluded that there was no obviously superior partitioning method, nor that providing the pKa values was consistently providing better predictions. However, the lipophilicity values provided by the Bayer tools (LogD and LogMA) did appear to give superior predictions compared to the other lipophilicity prediction sources. For this reason, we proceeded with the mean of those tools as the best lipophilicity prediction, as well as all partitioning and pKa prediction methods into the next round for the evaluation of clearance and Fu prediction tools (step 2).

### Step 2: Evaluation of Clearance and Fraction Unbound Predictions

After evaluating the physico-chemical properties determining passive distribution, we continued with the tools predicting key parameters depending on organism biology, specifically the Fu and clearance of compounds. We used the same IV PK data for validation as in the first step, but only using the best physico-chemical predictions as determined previously, while this time varying the Fu and clearance predictions.

For the prediction of the Fu, we had obtained values from seven *in silico* tools, as well as *in vitro* measured benchmark values. As expected, we found that the importance of Fu predictions depended on the clearance prediction approach used. When using *in vivo* plasma clearance benchmark values for model parameterisation, only marginal differences between the performances of the different Fu prediction tools were observed (SI-Fig. 7). But when predicting *in vivo* clearance using *in vitro* measured hepatic CLint values, we observed larger differences between the different Fu prediction tools (fig. 4AB). Our experimentally determined Fu values yielded better PK predictions than any *in silico* tool, which confirmed the validity of our benchmark reference values. However, the differences between the prediction qualities were overall relatively small. All Fu prediction tools led to Median Absolute Errors within the 2- to 3-fold range when using *in vitro* CLint values and there was no obvious systematic bias for under- or overprediction for any of the Fu prediction tools.

**Fig. 4:**
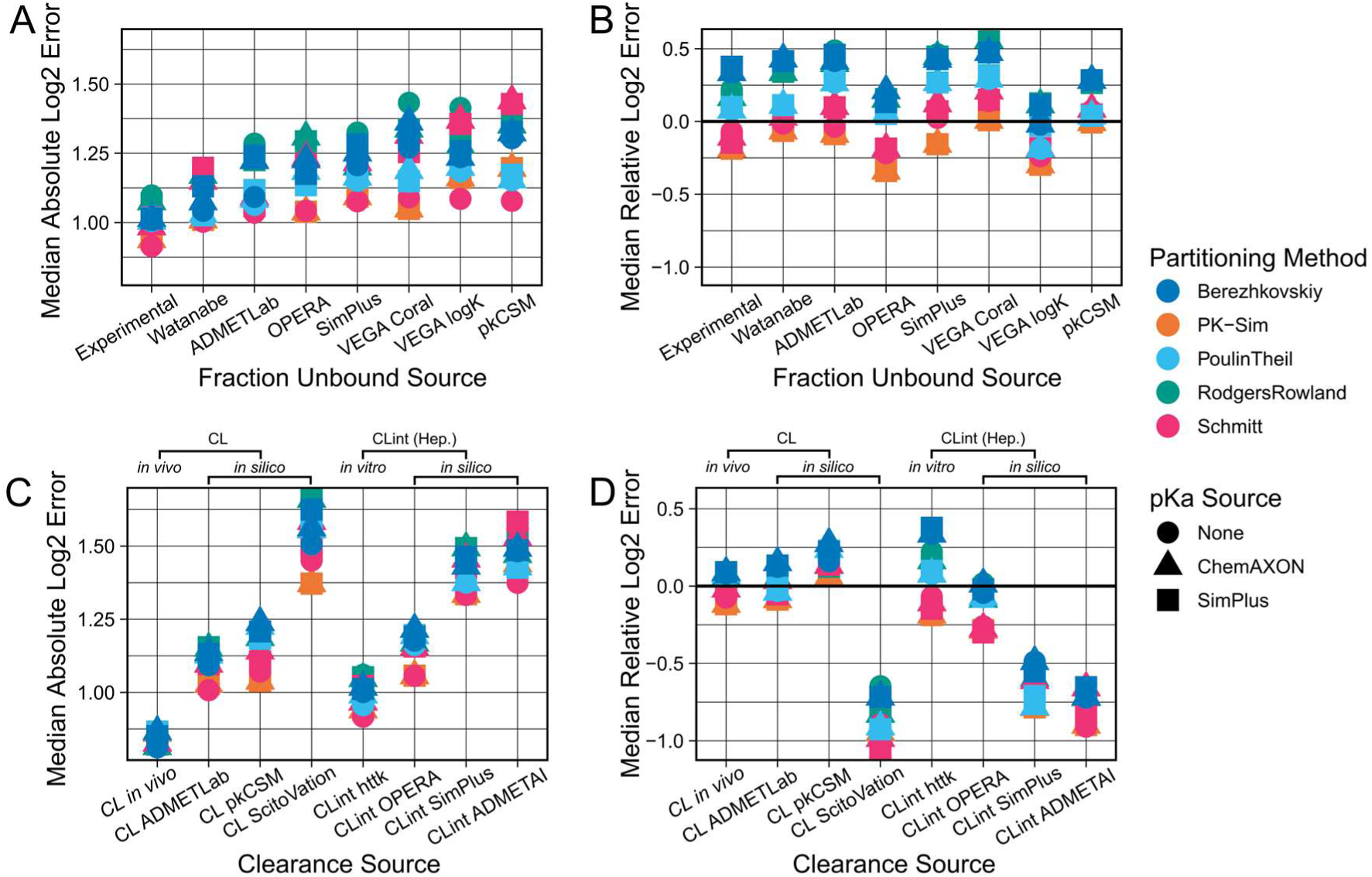
Comparison of predictive performances of different fraction unbound and clearance prediction sources (step 2). Combinations of all available parameterisation sources were evaluated against the collected IV dataset. Results shown were generated using benchmark values for parameterisation of the other parameters, i.e., *in vitro* CLint values from httk when comparing Fu predicting sources, and *in vitro* measured Fu values when comparing clearance predicting sources, as well as the mean of the two previously determined best lipophilicity prediction tools (LogD and LogMA Bayer) as lipophilicity values. The top row shows the Median Absolute Log2 Errors (A) and the Median Relative Log2 Errors (B) for different fraction unbound prediction methods. The bottom row shows the Median Absolute Log2 Errors (C) and the Median Relative Log2 Errors (D) for different clearance prediction methods. CL refers to plasma clearance values (measured or predicted), and CLint signifies hepatocyte intrinsic clearance (measured or predicted). *In vitro* measured CLint values from httk were only available for a subset of compounds (97 out of 143).

For the prediction of compound clearance, three *in silico* tools directly predicting plasma clearance, as well as our own previously used *in vivo* plasma clearance benchmark values, were available. Further, we had retrieved *in vitro* measured hepatic intrinsic clearance (CLint) values from httk, as well as *in silico* predictions of CLint values from OPERA, SimPlus and ADMETAI.

Before comparing the performances of the different clearance prediction strategies, we first evaluated whether activating passive renal excretion would improve or worsen PBK model simulations. Plasma clearance values already represent the total effect of all systemic clearance processes, so that adding passive renal clearance on top of them should theoretically lead to less accurate results. Whereas PBK models are expected to yield better PK predictions when passive renal excretion is incorporated if their *in vivo* clearance prediction is scaled up from hepatocyte-derived CLint values.

Overall, our results were consistent with these expectations. When adding passive renal clearance on top of the *in vivo* observed plasma clearance, prediction quality became worse and shifted from no bias to underprediction, whereas *in vitro* hepatocyte-based clearances improved from stronger to weaker overprediction of the PK data (SI-Fig. 8). For *in silico* predicted clearances, the situation was less straightforward. For instance, *in silico* CLint values predicted by SimPlus and ADMETAI already led to underpredictions of PK, which was then further exacerbated by additionally adding passive renal clearance. However, given the theoretical considerations, we continued our simulations by adding passive renal clearance on top of hepatocyte-scaled CLint, but not plasma clearance values.

Similar to the Fu, the *in vivo* observed plasma clearance benchmark values were the best input source for PBK model parameterisation (fig. 4C). All clearance prediction strategies yielded profoundly worse results than the benchmark *in vivo* clearance-based strategy, and almost all of them gave Median Absolute Errors worse than the 2-fold range. However, the differences between the clearance prediction tools were much larger than those between the Fu prediction tools. Out of the *in silico* predicted plasma clearance tools we found that ADMETLab gave the best predictions, followed by pkCSM and ScitoVation. While ADMETLab and pkCSM plasma clearance predictions led to a slight overprediction of the PK data, ScitoVation plasma clearance values led to a severe underprediction. We further confirmed these findings by directly comparing our *in vivo* plasma clearance values to the *in silico* predicted values of the different tools (SI-Fig. 9), which showed that ADMETLab’s plasma clearance values correlated best with our *in vivo* measured benchmark values.

*In vitro* hepatocyte CLint values from httk were the second-best clearance prediction source, after our *in vivo* plasma clearance benchmark values. Similar to what was observed for the *in silico* tools predicting plasma clearance values, the values from *in silico* CLint prediction tools also resulted in substantially worse PK predictions than the *in vitro* benchmark values. The best-performing CLint prediction tool was OPERA, followed by SimPlus and ADMETAI. However, we noted that for some compounds OPERA provided CLint values identical to the *in vitro* CLint values retrieved from httk (SI-Fig. 10). This suggested that those values were not true *in silico* predictions, which implies that the OPERA predictions may not be directly comparable to the other tools. Overall, when comparing *in silico* tools predicting plasma clearance values to the tools predicting hepatocyte CLint, most plasma clearance tools resulted in better PK predictions than most hepatocyte CLint prediction tools.

### Step 3: Evaluation of Solubility and Intestinal Permeability

In the third evaluation step, we investigated how to best predict parameters required for simulating oral administrations. In PK-Sim, these are primarily the solubility and intestinal permeability of a compound. We had obtained experimentally measured benchmark values of aqueous solubility, as well as predictions of aqueous solubility from four *in silico* tools (OPERA, ADMETLab, ProtoQSAR, SimPlus), and of Fasted State Simulated Intestinal Fluid (FaSSIF) and Fed State Simulated Intestinal Fluid (FeSSIF) solubility from SimPlus. For the intestinal permeability, no benchmark reference values were obtained. Instead, CACO2 permeability predictions from three *in silico* tools were used (OPERA, ADMETLab, ProtoQSAR), as well as MDCK permeability predictions from ADMETLab and SimPlus. Finally, we also obtained intestinal permeability predictions using the PK-Sim internal prediction equation, which is based on compounds’ molecular weight and lipophilicity (LogMA Bayer).

For the evaluation, we initially only used data from PK studies in which the administered formulation implied that the compound was already dissolved at administration (e.g., labelled as “solution” or “suspension”) and not in a solid state (e.g., “tablet” or “capsule”), since this additionally requires knowledge about the dissolution times of these formulations. We tested the mentioned prediction tools against the 286 concentration-time profiles (94 compounds) from those liquid formulation studies (fig. 5). However, no substantial difference was observed between the different parameterisation sources for either property. In the case of solubility, all *in silico* tools gave results similar to each other, and also comparable to the results of our experimentally measured benchmark values. Likewise, all intestinal permeability prediction tools gave comparable results.

**Fig. 5:**
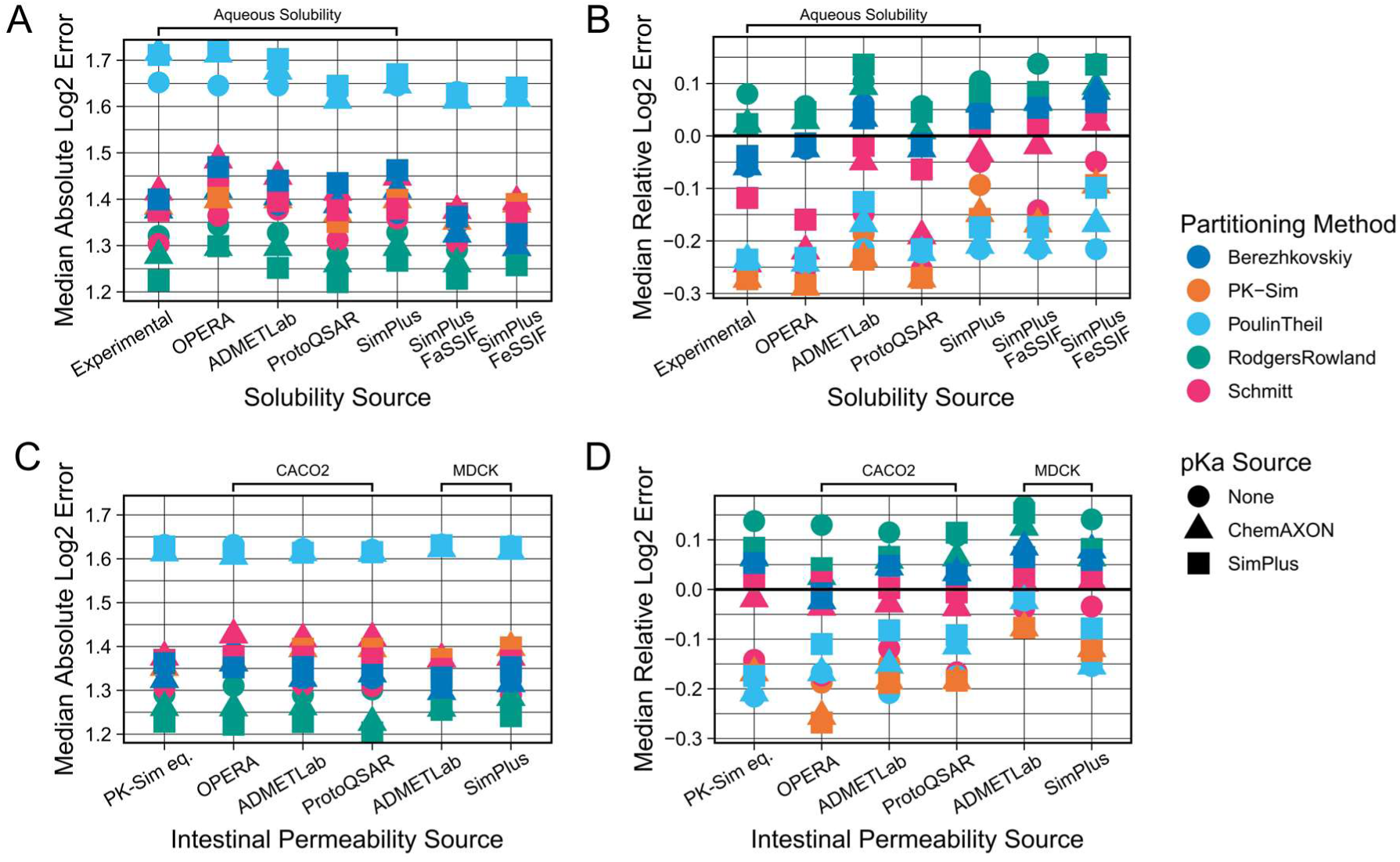
Comparison of predictive performances of different solubility and intestinal permeability prediction sources (step 3). Combinations of all available parameterisation sources were evaluated against the collected PO dataset (dissolved formulations). Results shown were generated using benchmark values for parameterisation of other parameters, i.e., *in vivo* plasma clearances, *in vitro* fraction unbound values and the mean of the two previously determined best lipophilicity prediction tools (LogD and LogMA Bayer) as lipophilicity values. The top row shows the Median Absolute Log2 Errors (A) and the Median Relative Log2 Errors (B) for different solubility prediction methods. The bottom row shows the Median Absolute Log2 Errors (C) and the Median Relative Log2 Errors (D) for different intestinal permeability prediction methods. For comparison of solubility predictions, the intestinal permeability values used were the PK-Sim internal equation values (PK-Sim eq.). For comparison of intestinal permeability sources, the solubility values used were SimPlus FaSSIF values. Results for the PK-Sim internal equation were generated with the Bayer LogMA predictions. Intestinal permeability predictions are either CACO2 or MDCK permeability predictions.

Further, we observed a general trend for overprediction of the velocity of oral absorption which resulted in a consistently strong underprediction of Tmax values, and a slight tendency for overprediction of Cmax values (SI-Fig. 11). We hypothesised that this might be because the *in silico* tools do not predict the PK-Sim specific intestinal permeability directly but instead were trained to predict *in vitro* measured CACO2 or MDCK permeabilities. However, when using such *in vitro* measured permeabilities, the standard procedure would be to scale these values, for example by using reference compounds, to the PK-Sim intestinal permeability parameter. Only when no measurements for reference compounds exist would one use the *in vitro* measured permeability values directly without scaling.

To take this into account, we extracted fitted PK-Sim intestinal permeability values of 56 compounds from Willmann et al. (2004) and then, for every *in silico* tool, determined a scaling factor based on the relationship between the values predicted by every tool and the assumed to be optimal values. While we found that there was a clear trend for the CACO2 values to be larger than the optimal PK-Sim intestinal permeability values (SI-Fig. 12), incorporating this scaling did not substantially improve PK predictions overall (SI-Fig. 13). Even though, it did reduce the strength of the bias in the underprediction of Tmax values.

Finally, we evaluated whether our conclusions based on the 94 compounds from liquid formulation studies would also hold true for the compounds of which we only had data from solid formulation studies. The simulation of these formulations required at least one additional parameter to describe the dissolution velocity of the solid formulations, which in reality will vary between different formulations. To at least determine which values might be appropriate average values, we tested different Lint80 dissolution times (10 – 30 min for capsules, 15 – 60 min for tablets) and then compared which average dissolution time would yield errors similar to what we had observed for the liquid formulations. Based on this we decided to use 25 min for formulations labelled “capsules” and 40 min for “tablets”, which extended the oral dataset for evaluation to 1200 PO concentration-time profiles (161 compounds). Using this larger dataset, all previously outlined conclusions were confirmed.

### Predictive Performances of Full HT-PBK Strategies

After evaluating step-by-step how to best predict every compound property required for HT-PBK modelling, we eventually assessed how well different types of HT-PBK strategies would predict the collected PK data overall. We identified the best strategies out of three classes. 1) As a benchmark comparison, we determined the performance of the best strategy overall, using *in vivo* and *in vitro* determined benchmark values of plasma clearance and Fu. 2) Additionally, the best fully *in silico*-based strategy was identified, for which we also considered property predictions coming from proprietary tools. 3) And finally, the best *in silico* strategy based exclusively on freely available tools was determined. The respective parameterisation strategies and their performances are presented in Table 1. Unsurprisingly, we found the strategy using benchmark reference values to be the most predictive. However, even fully *in silico*-based strategies yielded acceptable predictivity with 87%, or 89%, of Cmax values being predicted within 10-fold when using proprietary, or freely available prediction tools, respectively. Even more importantly, due to overestimation of the velocity of oral absorption in all strategies, the Cmax mispredictions outside the 10-fold range were mostly over-not underpredictions and therefore would lead to conservative, health-protective risk assessment conclusions. The performance of the best *in silico*-based HT-PBK approach is presented in fig. 6.

**Fig. 6:**
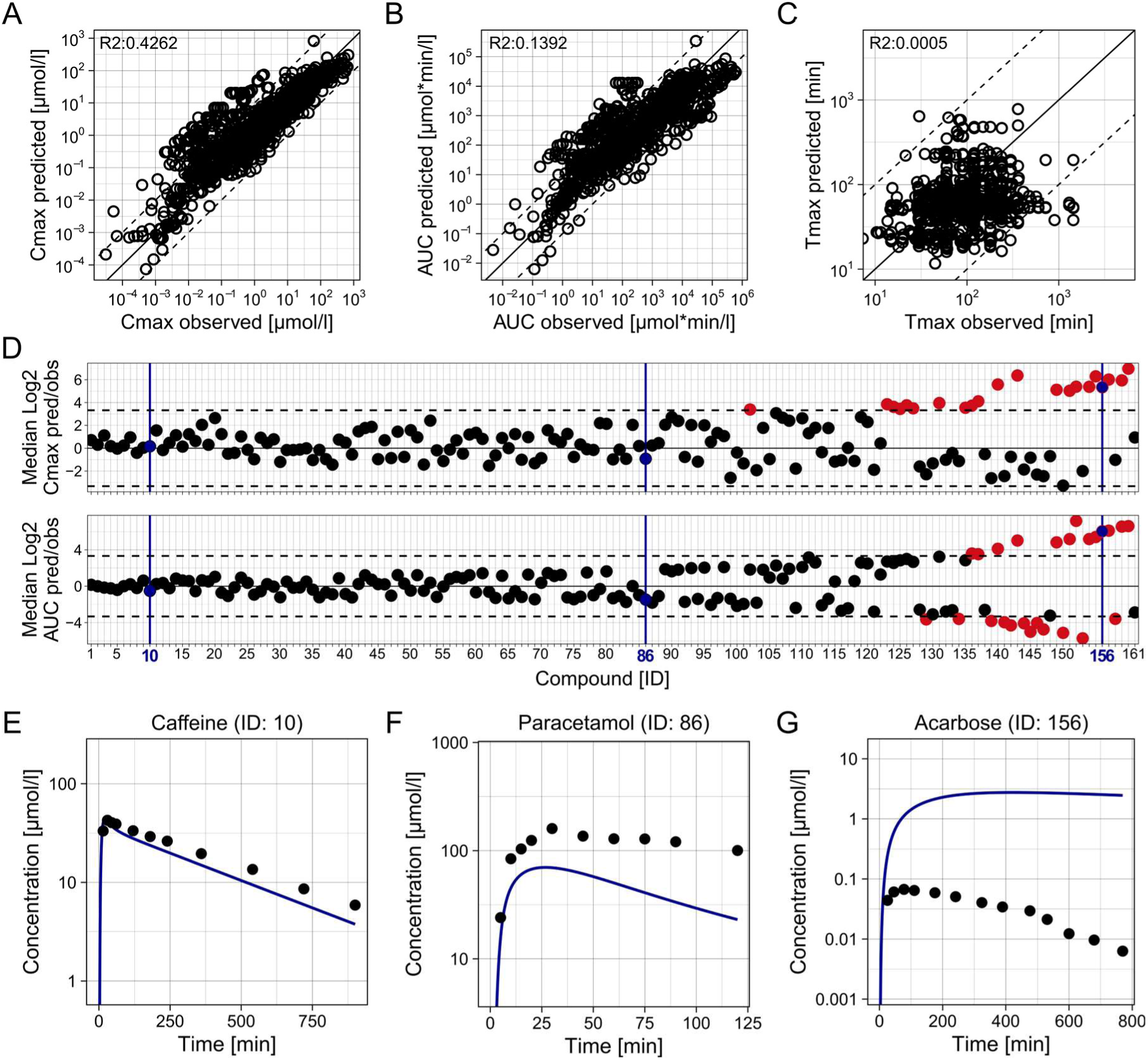
Predictive performance of the best fully *in silico*-based HT-PBK modelling strategy (proprietary). HT-PBK models were generated following the approach outlined in Table 1, then model simulations were compared against the collected PO dataset of 161 compounds (dissolved and solid formulations). (A) Predicted against observed Cmax values of all concentration-time profiles in the PO dataset. (B) Predicted against observed AUC values. (C) Predicted against observed Tmax values. Mind that A-C show individual concentration-time profiles with some compounds only represented by a single and others by multiple PK profiles. (D) The Median Log2 Cmax and AUC predicted/observed values of each compound in the PO dataset. Dashed lines indicate 10-fold errors. (E-G) Representative concentration-time profile predictions, and corresponding *in vivo* PK data, of three compounds embodying different levels of prediction quality. Blue colour marks the same representative compounds in D.

## Discussion

We here assembled a large dataset of healthy human *in vivo* concentration-time profiles (200+ compounds), as well as *in vitro* and *in silico* generated property predictions from various sources required for PBK modelling of the corresponding compounds. We systematically compared all possible HT-PBK modelling strategies to understand which prediction tools, and combinations thereof, perform best for parameterising PK-Sim to predict concentration-time data. Thereby, we quantified the expected accuracy of such HT-PBK predictions for typical pharmaceutical compounds.

For some input properties, especially lipophilicity and clearance, we found major differences in PBK model performances, while for other properties there was little variation. This may be due to larger differences in the predictive performances of the respective tools or due to differences in the sensitivity of the PBK model towards the different input parameters. Based on this observation, we conclude that generating and comparing PBK model simulations with different prediction tools for these critical parameters might be required to achieve more robust PBK model predictions. In contrast, for less sensitive parameters that may be of less importance. This could either be achieved by averaging property predictions of different tools, or by simulating ensembles of different PBK model variants. The latter would further allow to generate a distribution of simulation outcomes, thereby providing a confidence interval around simulation predictions. Such an explicit representation of uncertainty could then, for example, also be useful for conducting probabilistic risk assessment.

We also performed a large-scale comparison of the performances of frequently for PBK modelling used partitioning methods and found that all methods implemented in PK-Sim performed similarly well when based on high-quality lipophilicity values. However, for very high lipophilicity values, Poulin & Theil and Berezhkovskiy resulted in better PK predictions than other methods. It is likely though that more subtle differences in the performances of the different partitioning methods were not detectable with our approach, due to the intrinsic variability of our heterogeneous PK dataset gathered from several databases and the literature. Of note, even different PK profiles of the same compound but from different studies were not exactly identical to each other, which implies that our dataset contains intrinsic variability that even an ideal generic PBK model could never capture entirely. Consequently, this may have been the reason why we were unable to detect more subtle performance differences between the partitioning methods when using the highest quality lipophilicity values.

Besides their absolute errors, we also investigated whether any PK prediction strategies had biases for over- or underprediction. Our most noteworthy observation, apart from some tools having their individual biases, was that all HT-PBK modelling strategies seemed to overestimate the velocity of oral absorption, leading to an overprediction of Cmax and an underprediction of Tmax values. It is not fully clear why this occurred, as many factors influence oral absorption *in vivo.* Some of our fundamental assumptions, such as that “suspension” formulations were fully dissolved at administration, may have been inaccurate. Further, we parameterised our PBK models with a simple passive intestinal permeability parameter, thereby ignoring gut efflux transporters which are known to sometimes have profound impacts on the intestinal absorption of compounds. Ignoring such effects may have led to more rapid and full oral absorption in our simulations than occurred *in vivo*.

CACO2 or MDCK permeability values should, theoretically, account for such transporter effects (Volpe 2011). However, to be used in PK-Sim, they require scaling. Interestingly, our attempt to perform such a scaling using previously published fitted intestinal permeability values (Willmann et al. 2004) did indeed correct the biases in the underprediction of Tmax values, but the absolute PK prediction’s errors did not improve. One possible explanation for this result is that underpredictions of the velocity of oral absorption may have had a more severe impact on PK predictions than its overprediction. And for this reason, we observed lower Absolute Log2 Errors when using CACO2 values directly, despite scaled CACO2 values being potentially more correct representations of *in vivo* oral permeability. For applications in the pharmaceutical domain, it may therefore be of interest to further investigate such CACO2 scaling approaches. However, for toxicological applications, an overprediction of the velocity of oral absorption may even be preferable since it leads to a health-protective bias in risk assessments.

Similar to gut transporters, we were also unable to explicitly account for the contribution of transporter effects in other body tissues that may potentially alter compounds’ PK. Performing high-throughput assessments requires a generic modelling approach and readily available, homogeneous input data from the same sources to be able to systematically compare the predictive performances of strategies. Unfortunately, such data were not available for transporters. Further, our approach solely focused on parent compounds and neglected the issues of bioactivation and metabolism, since the formation of metabolites cannot be predicted quantitatively. Also, it would be valuable to gain more insight into which compound properties (physico-chemical properties, clearance pathways, transporter affinity etc.) are correlated with lower or higher prediction accuracies, since we do observe large differences between prediction accuracies among different compounds.

Because most compounds in our dataset were typical pharmaceutical compounds, it is further likely that many of them were present in the training data of the *in silico* tools we evaluated, potentially biasing our analysis. This was most evident for the CLint values gathered from OPERA, which, for some compounds, were identical to the *in vitro* CLint values from httk. It might be insightful to analyse in more detail how prediction accuracies of *in silico* tools differ between those compounds on which the models were trained originally and those compounds that were outside their training datasets. Nevertheless, from a practical point of view, the fact that some *in silico* tools may be able to recall data from larger or higher quality training datasets may also be interpreted as a strength of these tools.

Eventually, we showed that it is possible to generate reliable HT-PBK models for the prediction of IV and PO PK of pharmaceuticals. In principle, it is also possible to apply our approach to other classes of compounds, although the validation of this is hampered by the absence of comparable concentration-time data for validation. Potential applications of such HT-PBK modelling strategies are vast, and there have been many efforts recently in both pharmacology and toxicology to establish such strategies for different use cases and based on different approaches. Many of them, however, relied exclusively on rodent data for their validation (Schneckener et al. 2019; Kamiya et al. 2021; Naga et al. 2022; Punt et al. 2022b; Obrezanova et al. 2022; Mavroudis et al. 2023; Handa et al. 2023; Führer et al. 2024). Others did perform predictions for humans but relied in their validation of prediction quality on summary PK parameters, like Cmax or AUC values (Punt et al. 2022a; Miljković et al. 2021; Li et al. 2023). The problem with such approaches is that they do not consider the full information about the quality of the predicted concentration-time curve as a whole. Hence, they are unable to deconvolute the biases of individual input sources, that can potentially compensate each other, and which may obscure inaccuracies in model parameterisation. This is why we here relied on full concentration-time profiles, and multiple summary parameters, for evaluation, as it would be done by an expert for the traditional development of a PBK model.

Another now frequently used strategy is to use ML and AI techniques to directly predict summary PK parameters, like the Cmax, AUC or bioavailability of compounds (Schneckener et al. 2019; Miljković et al. 2021; Fagerholm et al. 2021; Obrezanova et al. 2022). However, so far, this approach did not perform as well as using *in silico* predicted properties to then inform mechanistic simulations based on PBK models. For this reason, we here followed the latter strategy of using ML models to predict mechanistically meaningful compound properties to then input these into the PK-Sim PBK model, which incorporates *ab initio* expert knowledge about known body physiology into our high-throughput predictions. Besides yielding more accurate predictions of PK parameters, this approach maintains the explainability of our models and their predictions. Since every PBK model parameter has a physiological meaning, the approach further enables investigation into why certain compounds might be mispredicted by different strategies and to understand which property mispredictions are responsible for inaccurate PK predictions. Furthermore, this approach makes it possible to leverage PBK models’ ability to mechanistically extrapolate predictions, for example, to special populations or other exposure scenarios.

Finally, the here presented HT-PBK modelling is a promising tool for applications in both pharmacology and toxicology. In drug discovery, HT-PBK models have the potential to aid rapid compound selection and optimisation decisions. While HT-PBK modelling may not initially match the accuracy of traditional PBK models, it can provide a base model which may then be progressively refined throughout the drug development cycle to meet escalating demands for accuracy and detail. For toxicological risk assessment, accurate predictions of internal organ concentrations are key to replace animal testing with *in vitro* and *in silico* methods. Such predictions can, for example, assist with the prioritisation and classification of chemicals, or provide valuable information for quantitative *in vitro* to *in vivo* extrapolation. For a new methodology like HT-PBK modelling to be used in Next Generation Risk Assessment, however, it is key to validate its predictive performance, in order to generate the high confidence required for its regulatory use. Using a large, heterogeneous PK dataset, we here showed that the outlined HT-PBK modelling strategies are fit-for-purpose for such applications.

## Acknowledgements

We thank Susana Proença, Nynke Kramer and Huan Yang for discussions about the simulation analysis strategy and Pavel Balazki for advice on the technical implementation of PK-Sim model simulations in R.

## Funding

This work was performed in the context of the ONTOX project (https://ontoxproject.eu/) that has received funding from the European Union’s Horizon 2020 Research and Innovation programme under grant agreement No 963845. ONTOX is part of the ASPIS project cluster (https://aspiscluster.eu/).

## Ethical standards

This manuscript exclusively relies on already publicly available clinical data.

## Conflicts of interest

Marina García de Lomana is an employee of Bayer AG. Rita Ortega-Vallbona and Eva Serrano-Candelas are employees of ProtoQSAR SL. Alicia Paini was an employee of esqLABS GmbH when this work was conceived. Stephan Schaller is founder and managing director of esqLABS GmbH. All authors declare that they have no conflict of interest.

## Abbreviations

ADME: Absorption, Distribution, Metabolism, and Excretion
AI: Artificial Intelligence
AUC: Area under the curve
CL: Clearance
CLint: Intrinsic hepatic clearance
Cmax: Maximum concentration
HT-PBK: High-throughput
PBK IV: Intravenous
LogD: Logarithm of distribution coefficient
LogMA: Logarithm of membrane affinity partition coefficient
LogP: Logarithm of octanol-water partition coefficient
ML: Machine Learning
Tmax: Time to maximum concentration
PK: Pharmacokinetics
TK: Toxicokinetics
PBK: Physiologically based kinetic
PBPK: Physiologically based pharmacokinetic
PBTK: Physiologically based toxicokinetic
PO: Peroral

## Supplementary Information

### SI-1: Extensive Description of the PK Data Retrieval Process

We downloaded all concentration-time data from the PK-DB, CvT-DB and OSP Observed Data Repository databases and filtered them for healthy, adult human subjects. We excluded all studies performed on special populations including paediatric (< 17 years), geriatric (> 60 years) or diseased populations, which we expected might display altered pharmacokinetics. Some selected conditions, like “migraine”, “psychiatric disorder”, “asthma”, “diabetes” and even “liver disease (minimal)” were permitted, however. But, for example, “liver disease (severe)” or “renal disease (end-stage)” were excluded. We filtered for profiles of administered parent compounds, discarding data of metabolites, and only selected those data that were measured in blood, plasma or serum, and only after single dose intravenous or oral administration. Further, we excluded all studies with concomitant administrations of other drugs to avoid data from drug-drug interaction studies, however, “control” or “placebo” groups from such studies were permitted. Also, we excluded endogenous compounds like ethanol or glucose. If information about both nominal dose and salt-free active pharmaceutical ingredient (API) was provided, we retrieved the salt-free API value as the dose value. When only a nominal dose was provided, we assumed this was referring to the salt-free API dose. Labels of formulations for oral administration (“tablet”, “capsule”, “solution” etc.) were extracted as provided.

After retrieving concentration-time profiles from above mentioned databases, we enriched our dataset by manually digitising more concentration-time data from the literature. For this, we followed the same strategy as before, aiming for data from healthy adults. However, in some cases, for example for certain chemotherapeutics, it proved difficult to find PK data for healthy adults or subjects younger than 60 years. Since we were able to inspect those studies manually, and to ensure that they were not performed on those special populations due to a suspected difference in pharmacokinetics, we occasionally decided to include them, if no other data on such compounds was available.

Next, we excluded data points which had a concentration value numerically close to zero (< 1e-7) after 4 h, since some studies possessed unreasonably low concentration values at the end of their concentration-time profiles, presumably representing the lower limit of quantification, or due to digitisation inaccuracies. Also, we excluded all concentration-time profiles with less than two time points. If we had retrieved more than 30 concentration-time profiles for a given compound and a given route of administration, then we excluded all profiles from individuals (where number of subjects = 1), as long as there were at least 20 other concentration-time profiles left. If the number of profiles was then still higher than 30, we excluded the studies with the least number of subjects until we had no more than 60 concentration-time profiles per study and route. Finally, we generated dose normalised plots for all compounds where multiple concentration-time curves were available to visually confirm that studies were consistent with each other and that there were no extreme outliers among the data. When we found noticeable differences between concentration-time profiles in the dose normalised plots, we looked up the corresponding original publications and either corrected or excluded those entries.

---

**SI-Fig. 1:**
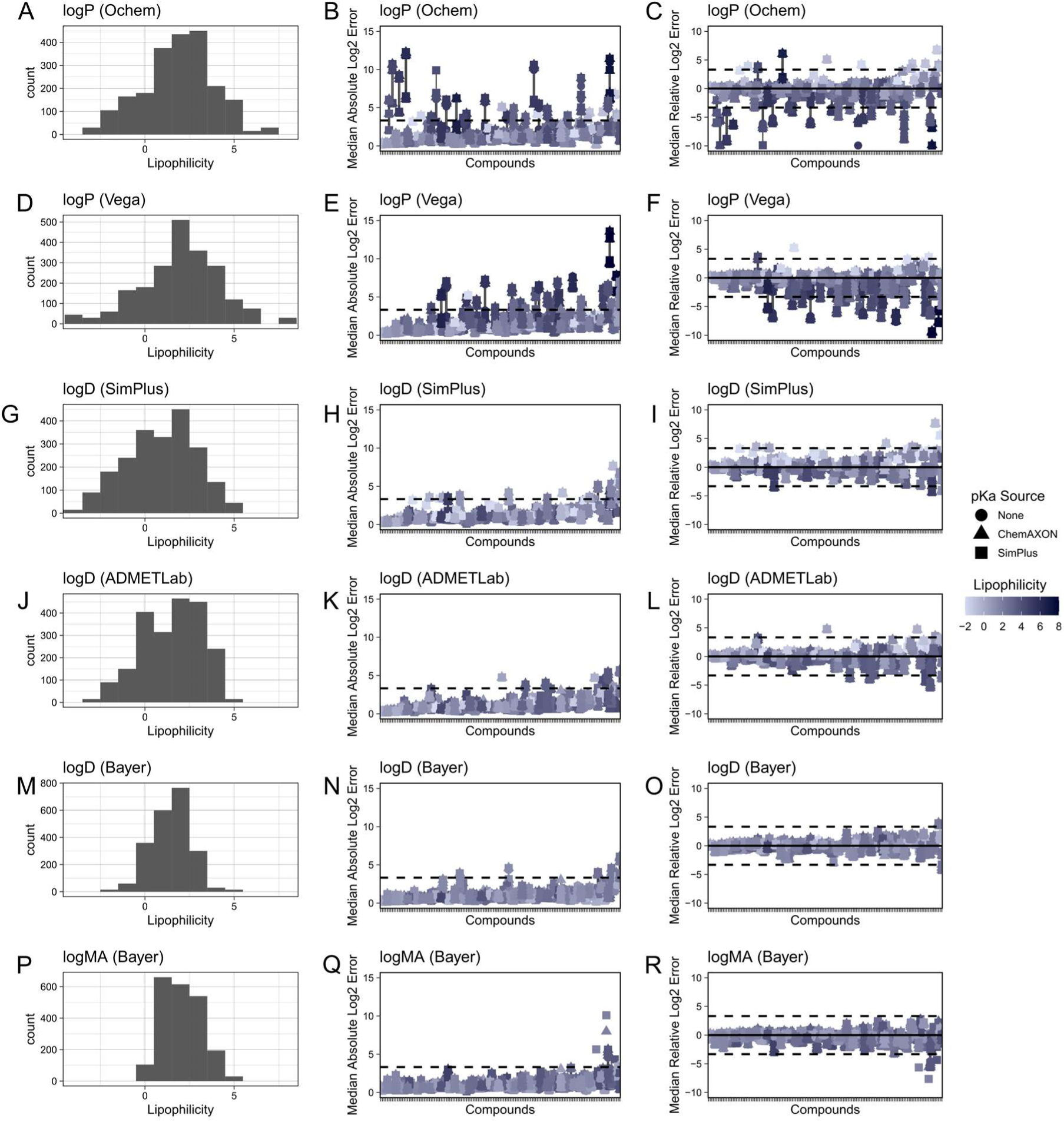
Comparison of predictive performances of different lipophilicity prediction tools coloured by lipophilicity value (step 1). Combinations of all available PBK model parameterisation sources were evaluated against the collected IV dataset. Clearance and fraction unbound were parameterised using *in vivo* and *in vitro* reference benchmark values, respectively. Left column shows histograms of predicted lipophilicity values by the various prediction tools. Middle column shows Median Absolute Log2 Errors of every compound, colour-coded by lipophilicity value predicted by the tool, and right column shows Median Relative Log2 Errors. Dashed lines indicate 10-fold errors.

**SI-Fig. 2:**
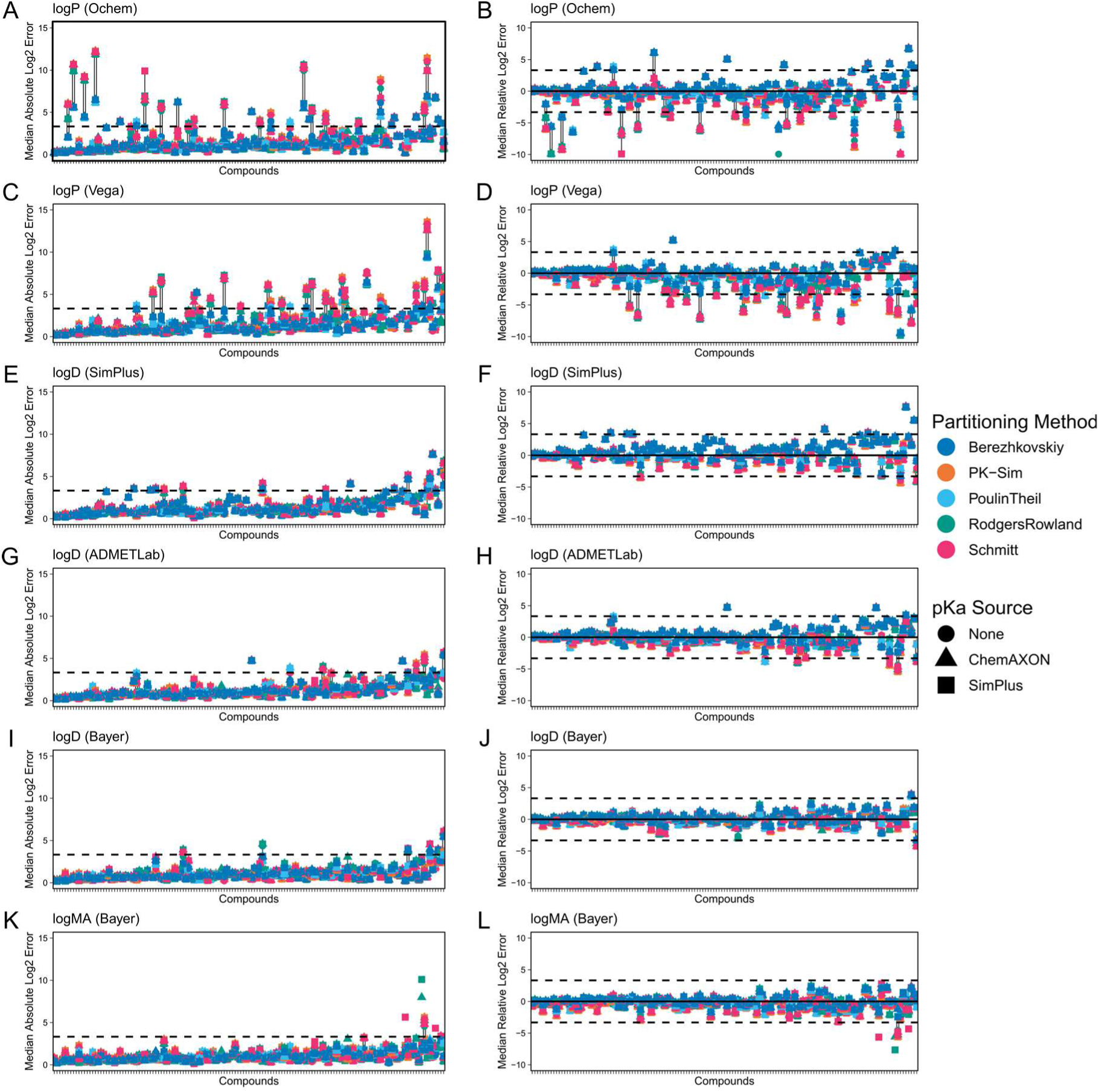
Comparison of predictive performances of different lipophilicity prediction tools coloured by partitioning method (step 1). Combinations of all available PBK model parameterisation sources were evaluated against the collected IV dataset. Clearance and fraction unbound were parameterised using *in vivo* and *in vitro* reference benchmark values, respectively. Left column Median Absolute Log2 Errors of every compound, colour coded by partitioning method used for simulation, and right column shows Median Relative Log2 Errors. Dashed lines indicate 10-fold errors.

**SI-Fig. 3:**
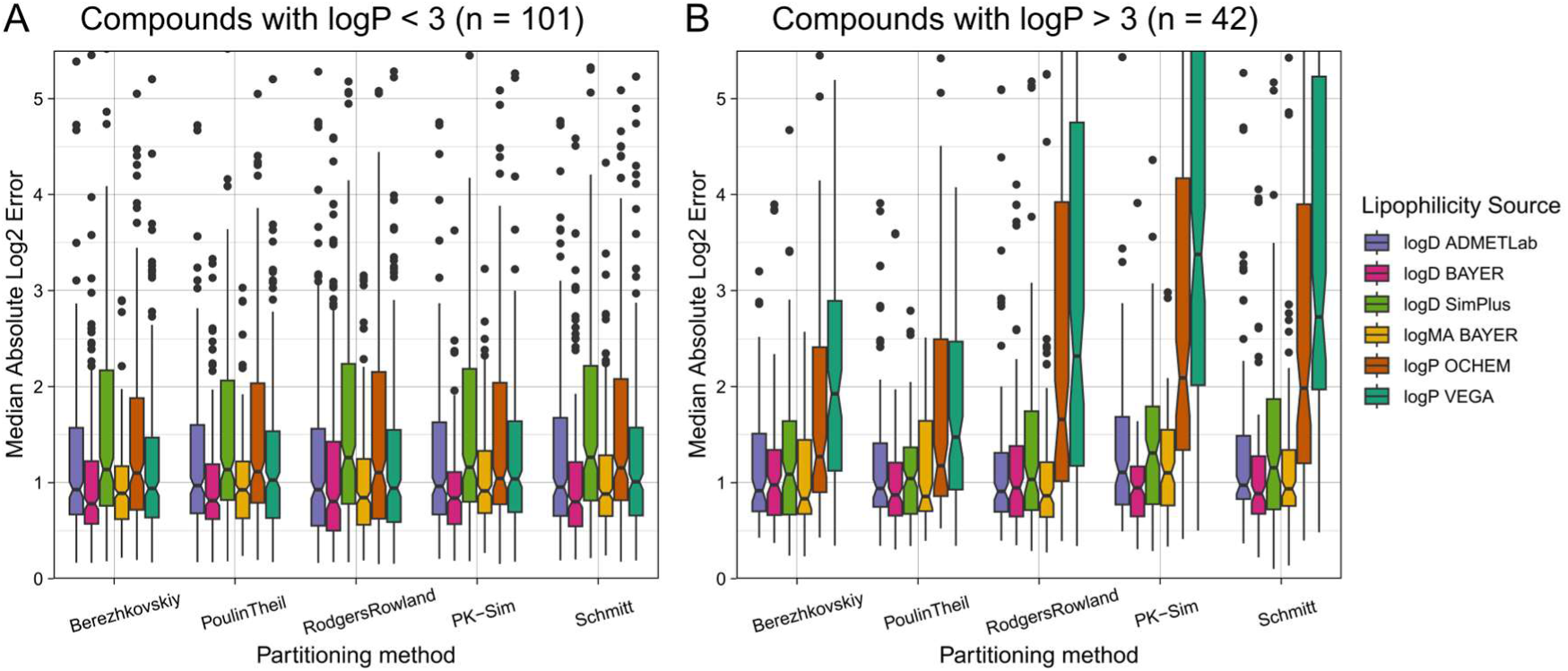
Comparison of predictive performances of lipophilicity prediction tools and partitioning methods separated by compound lipophilicity. Combinations of all available PBK model parameterisation sources were evaluated against the collected IV dataset. Clearance and fraction unbound were parameterised using *in vivo* and *in vitro* reference benchmark values, respectively. Lipophilicity values were set using the mentioned LogP and LogD *in silico* prediction tools. Plots show Median Absolute Log2 Errors as a measure of prediction precision. A shows results of compounds with LogP < 3 (n = 101), B shows results of compounds with LogP > 3 (n = 42).

**SI-Fig. 4:**
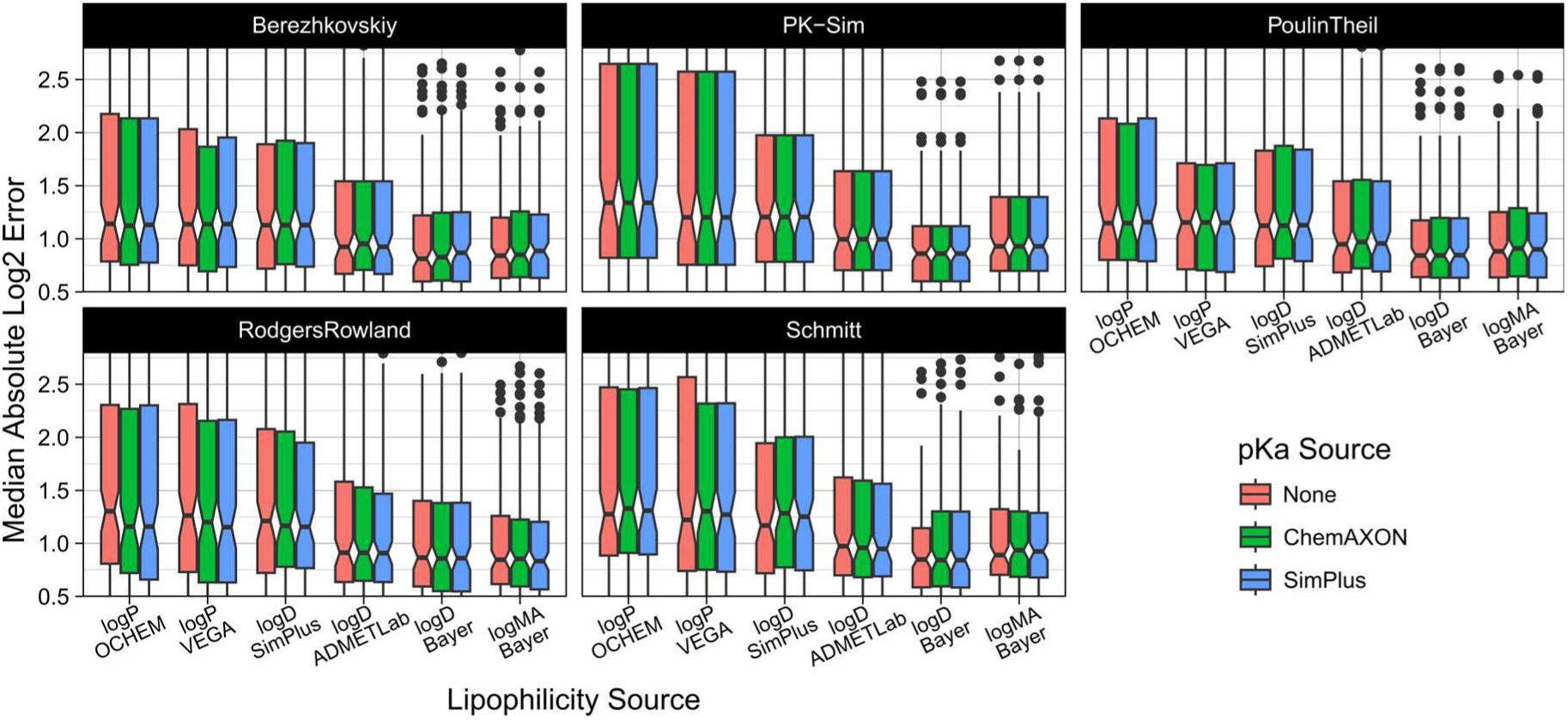
Comparison of predictive performances of pKa prediction tools. Combinations of all available PBK model parameterisation sources were evaluated against the collected IV dataset. Clearance and fraction unbound were parameterised using *in vivo* and *in vitro* reference benchmark values, respectively. Lipophilicity values were set using the mentioned LogP and LogD *in silico* prediction tools. Plots show Median Absolute Log2 Errors as a measure of prediction precision.

**SI-Fig. 5:**
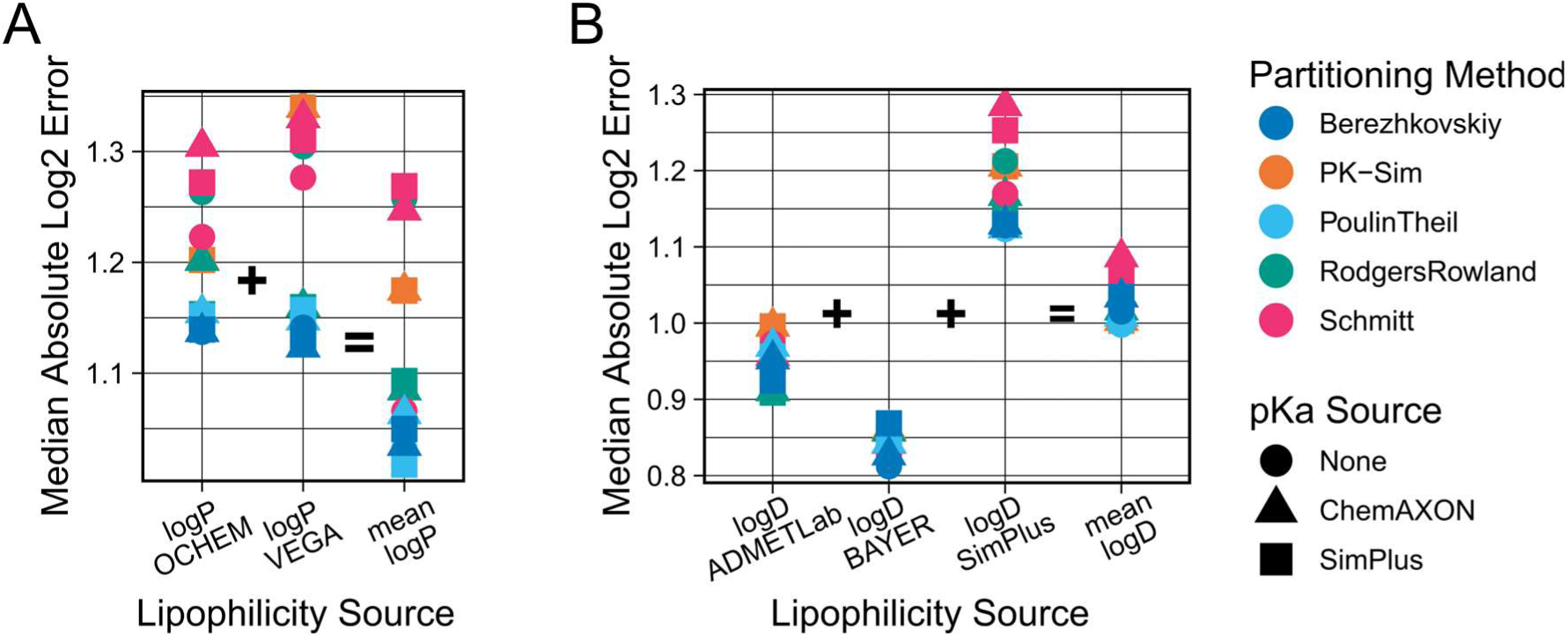
Predictive performances of consensus strategies for averaging lipophilicity predictions. Combinations of all available PBK model parameterisation sources were evaluated against the collected IV dataset. Clearance and fraction unbound were parameterised using *in vivo* and *in vitro* reference benchmark values, respectively. Lipophilicity values were set using the mentioned LogP and LogD *in silico* prediction tools, as well as the averages of those. Plus signs indicate which predictions of the individual LogP or LogD tools were combined to achieve the mean results marked by equal signs.

**SI-Fig. 6:**
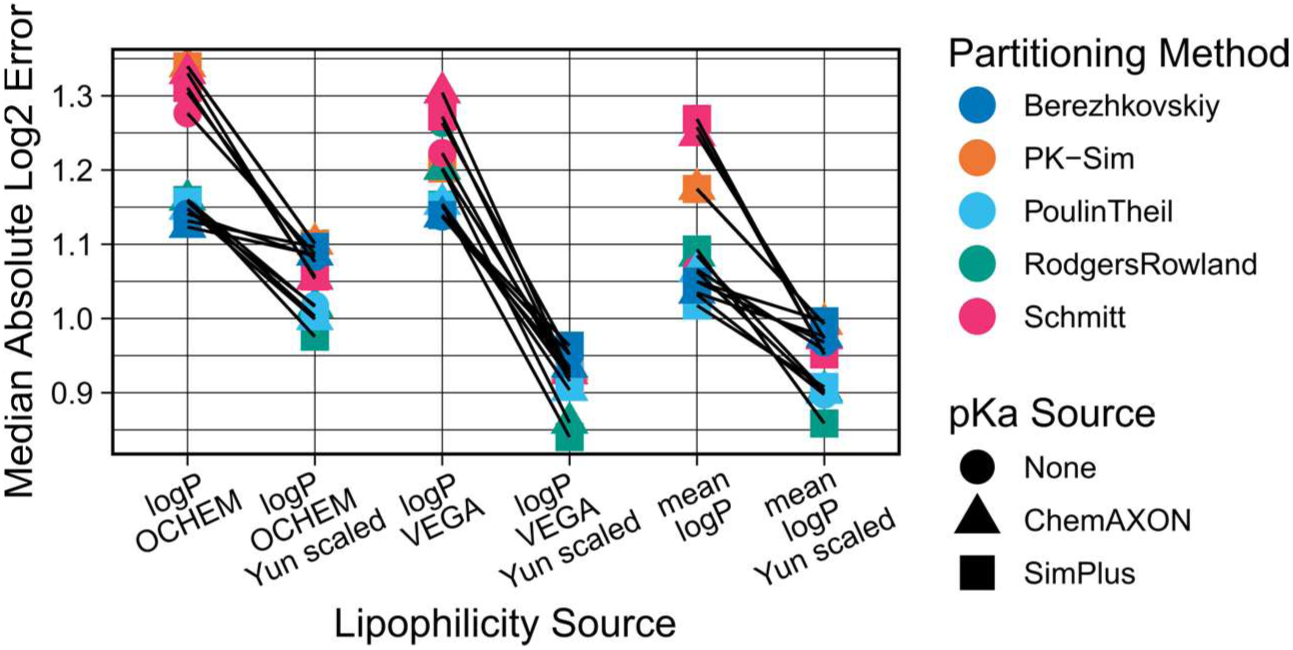
Change in predictive performance when using an empirical regression equation to convert LogP lipophilicity to LogMA. Combinations of all available PBK model parameterisation sources were evaluated against the collected IV dataset. Clearance and fraction unbound were parameterised using *in vivo* and *in vitro* reference benchmark values, respectively. Lipophilicity values were set using the mentioned LogP *in silico* prediction tools VEGA, OCHEM and their average. Additionally, those LogP values were converted to LogMA values by an empirical regression equation established by Yun et al. (2014).

**SI-Fig. 7:**
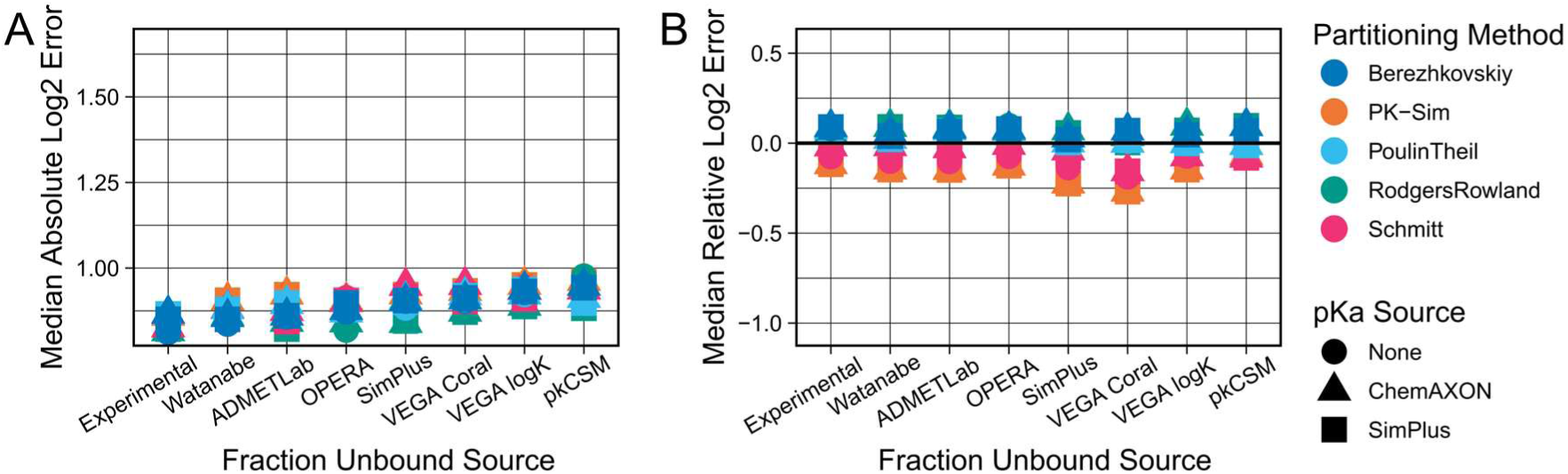
Comparison of Fu prediction tools when using benchmark *in vivo* plasma clearance values for PBK model parameterisation. Combinations of all available parameterisation sources were evaluated against the collected IV dataset. Results shown were generated using *in vivo* observed plasma clearance benchmark values, and the mean of the two previously determined best lipophilicity prediction tools (LogD and LogMA Bayer) as lipophilicity values. A: Median Absolute Log2 Errors as a measure of prediction precision. B: Median Relative Log2 Errors as a measure of systematic prediction bias.

**SI-Fig. 8:**
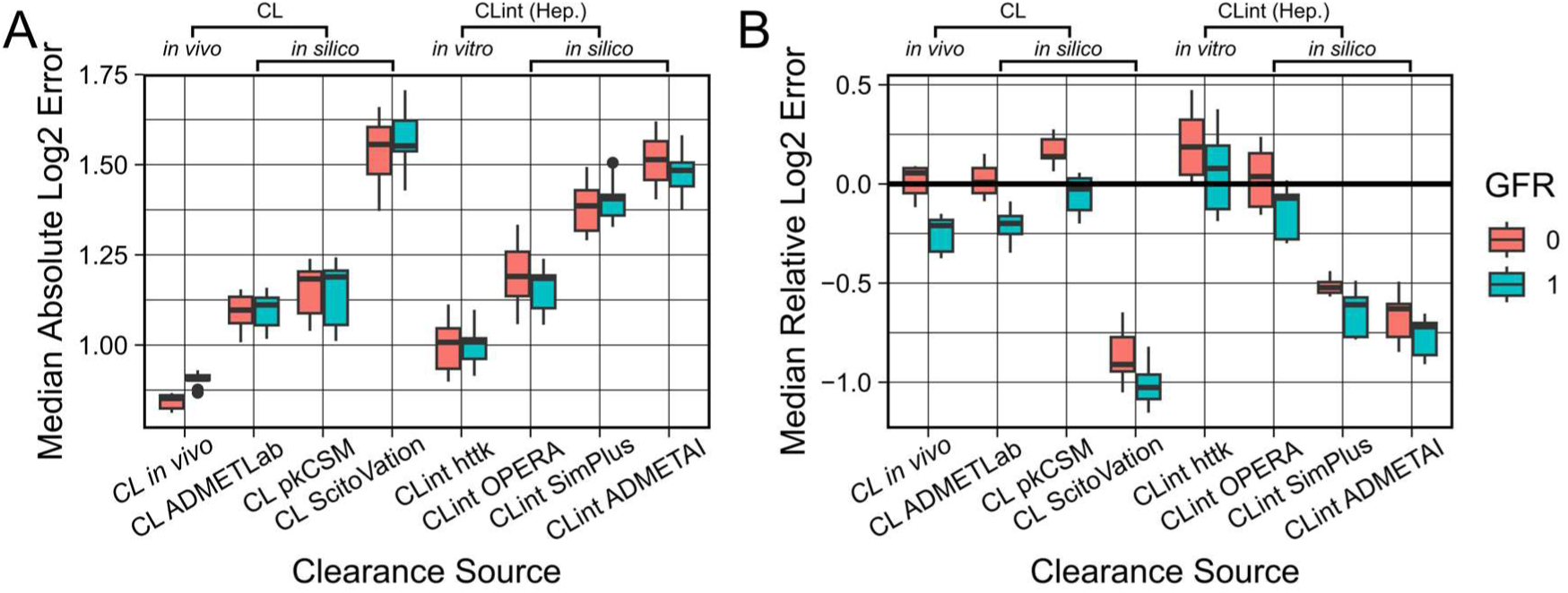
Effects of adding passive renal clearance on the predictive performances of HT-PBK predictions. Combinations of all available parameterisation sources were evaluated against the collected IV dataset. Results shown were generated using *in vitro* measured Fu benchmark values, and the mean of the two previously determined best lipophilicity prediction tools (LogD and LogMA Bayer) as lipophilicity values. GFR rate of 1 signifies adding passive renal clearance to the PBK model, and a value of 0 indicates leaving it out. CL refers to plasma clearance values (measured or predicted), CLint stands for hepatic intrinsic clearance (measured or predicted). A: Median Absolute Log2 Errors as a measure of prediction precision. B: Median Relative Log2 Errors as a measure of systematic prediction bias.

**SI-Fig. 9:**
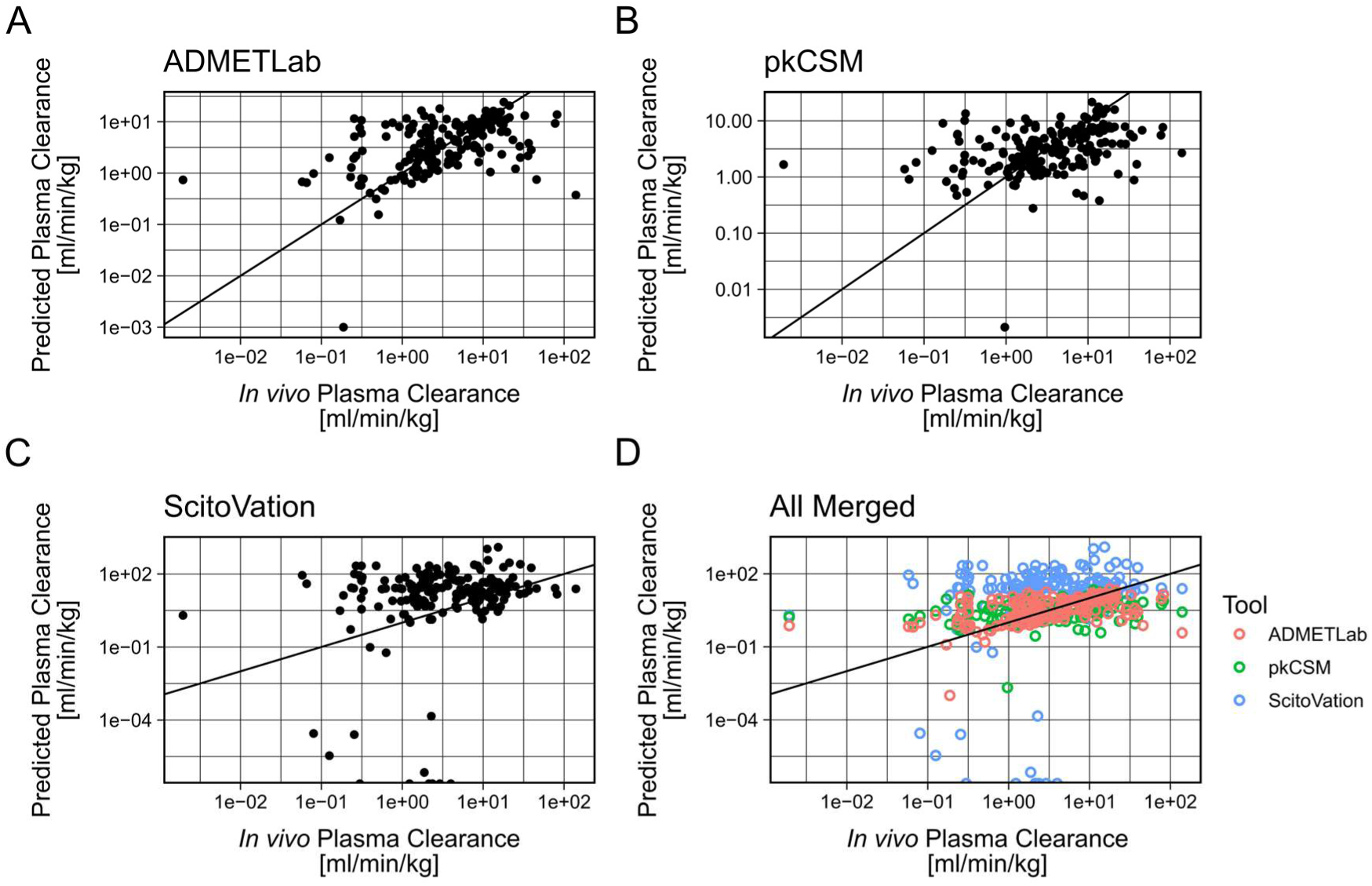
*In silico* predicted plasma clearance values from different tools against *in vivo* observed values. *In vivo* values were collected from the literature, *in silico* values were predicted using the corresponding tools. A-C: *In silico* predicted plasma clearance values from ADMETLab, pKCSM, and ScitoVation, respectively. D: All predicted values combined in a single plot.

**SI-Fig. 10:**
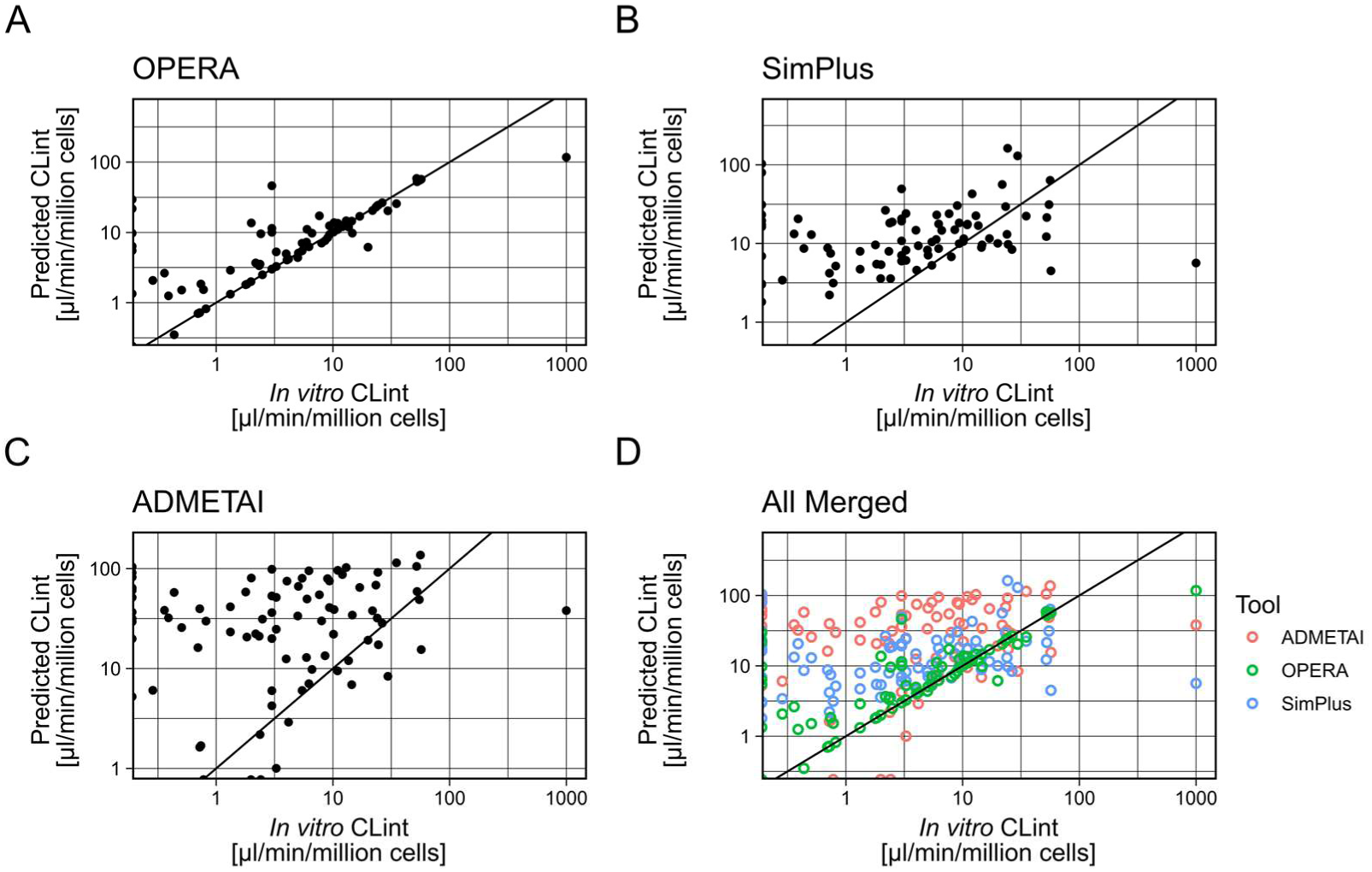
*In silico* predicted hepatocyte CLint values from different tools against *in vitro* measured values. *In vitro* CLint values were taken from httk version 2.2.1, *in silico* values were predicted using the corresponding tools. A-C: *In silico* predicted plasma clearance values from OPERA, SimPlus (ADMET Predictor), and ADMETAI, respectively. D: All predicted values combined in a single plot.

**SI-Fig. 11:**
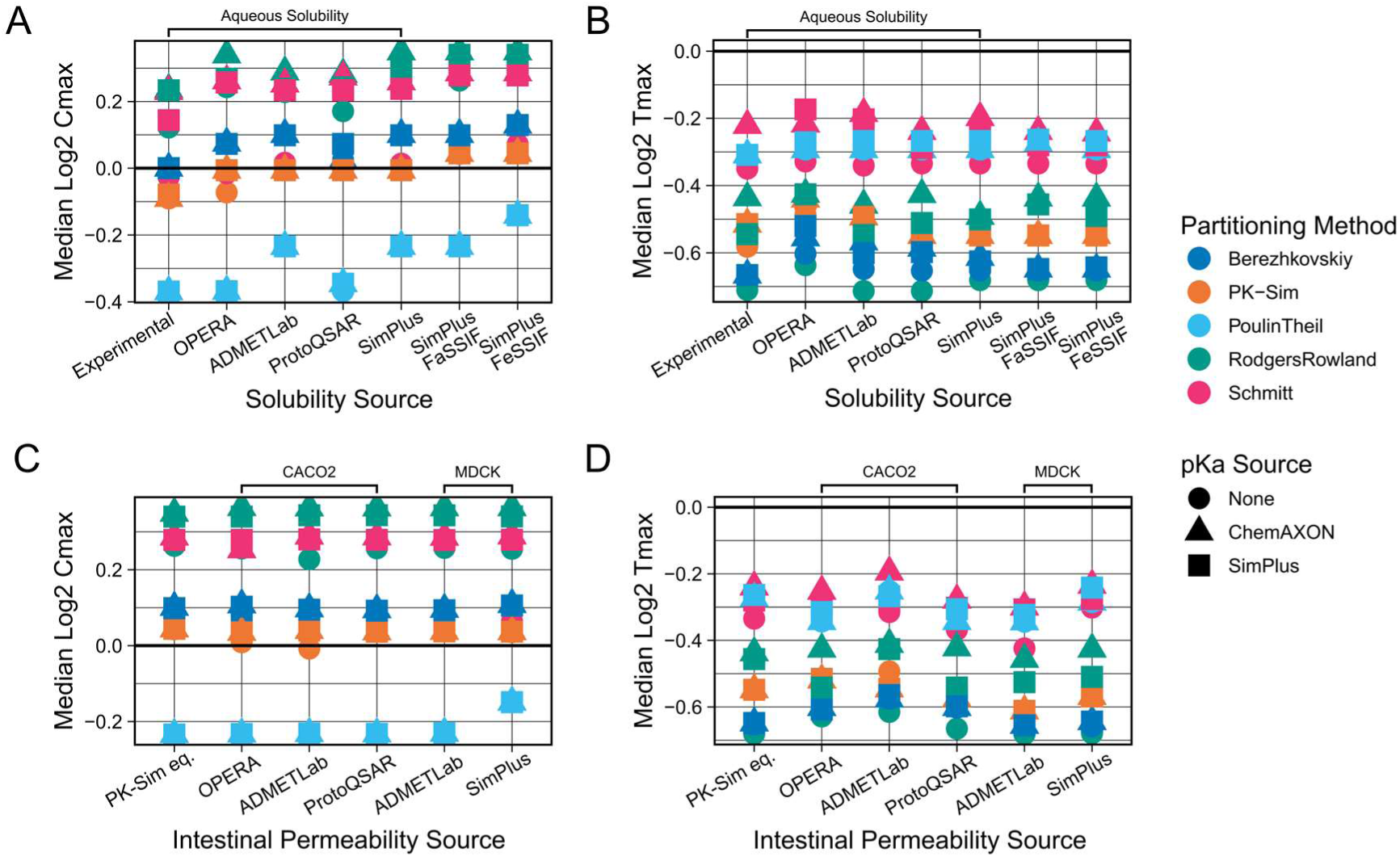
Cmax and Tmax prediction biases of different solubility and intestinal permeability prediction sources (step 3). Combinations of all available parameterisation sources were evaluated against the collected PO dataset (dissolved formulations). Results shown were generated using benchmark values for parameterisation of other parameters, i.e., *in vivo* plasma clearances, *in vitro* fraction unbound values and the mean of the two previously determined best lipophilicity prediction tools (LogD and LogMA Bayer) as lipophilicity values. The top row shows the Median Log2 Cmax predicted/observed (A) and the Median Log2 Tmax predicted/observed (B) for different solubility prediction methods. The bottom row shows the Median Log2 Cmax predicted/observed (C) and the Median Log2 Tmax predicted/observed (D) for different intestinal permeability prediction methods. For comparison of solubility sources, the intestinal permeability source used was the PK-Sim internal equation (PK-Sim eq.). For comparison of intestinal permeability sources, the solubility values used were the SimPlus FaSSIF values. Results for the PK-Sim internal equation were generated by inputting the Bayer LogMA predictions. Intestinal permeability predictions are either CACO2 or MDCK permeability predictions.

**SI-Fig. 12:**
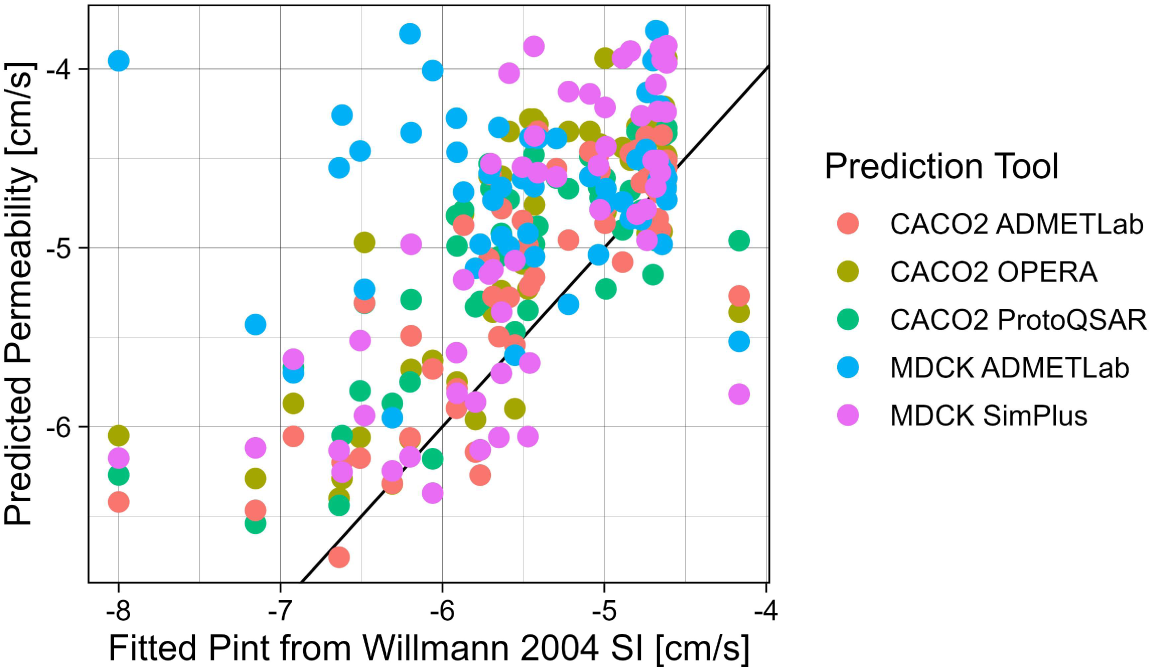
*In silico* predicted intestinal permeability values from different tools against optimal fitted Pint values. Optimal fitted Pint values were taken from Willmann et al. (2004).

**SI-Fig. 13:**
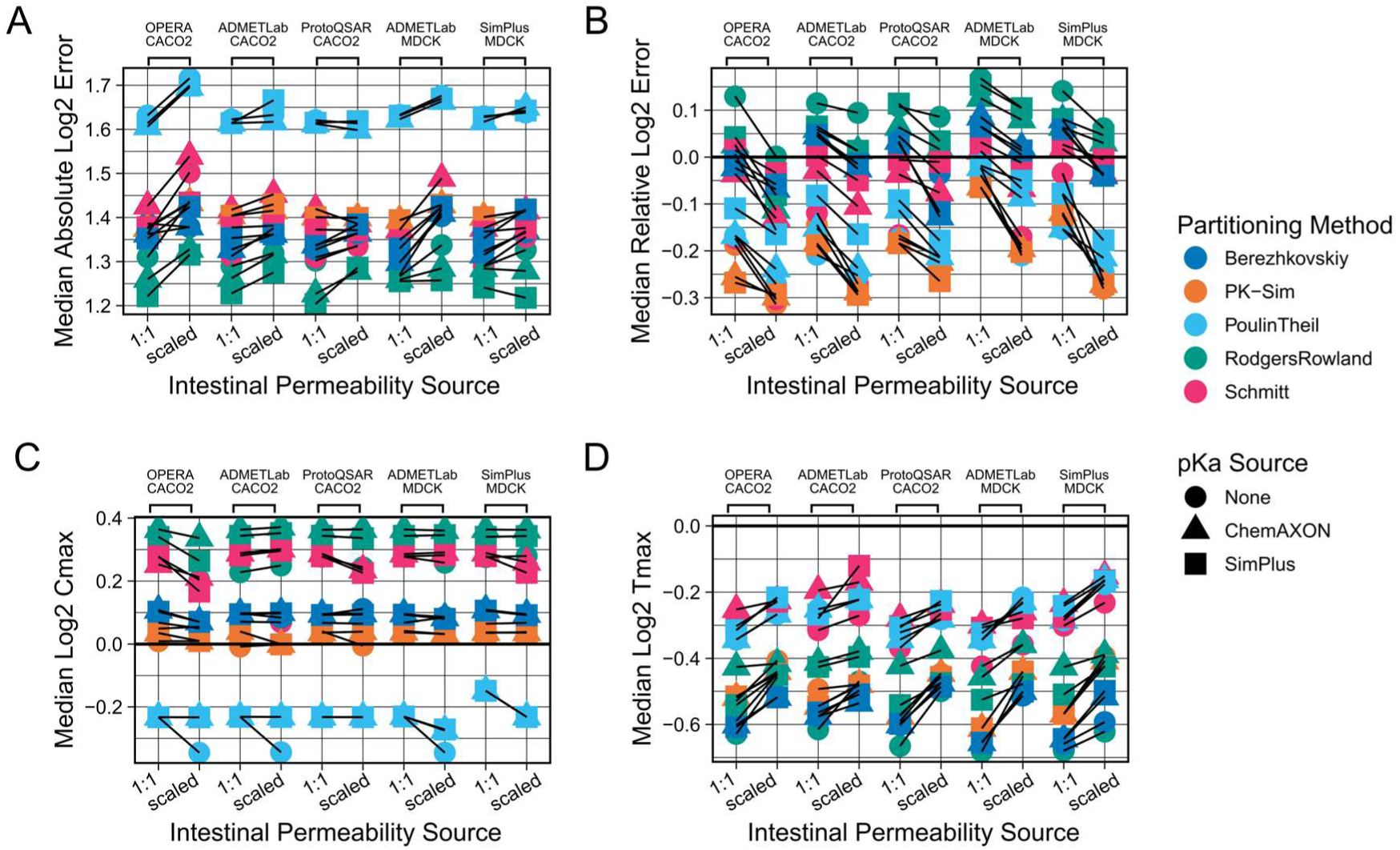
Effect of scaling intestinal permeability predictions on different performance metrics. Combinations of all available parameterisation sources were evaluated against the collected PO dataset (dissolved formulations). Results shown were generated using benchmark values for parameterisation of other parameters, i.e., *in vivo* plasma clearances, *in vitro* fraction unbound values and the mean of the two previously determined best lipophilicity prediction tools (LogD and LogMA Bayer) as lipophilicity values. SimPlus FaSSIF values were used as solubility values. The top row shows the Median Absolute Log2 Error (A) and the Median Relative Log2 Error (B) for different intestinal permeability prediction approaches. The bottom row shows the Median Log2 Cmax predicted/observed (C) and the Median Log2 Tmax predicted/observed (D). Lines connect the simulation results based on the same *in silico* prediction tool, either used directly (1:1) or after scaling using optimal intestinal permeability values from Willmann et al. (2004).

## Notes

### Competing Interest Statement

The authors have declared no competing interest.

